# HEXOKINASE1 interferes with cytokinin synthesis and strigolactone perception during sugar-induced shoot branching

**DOI:** 10.1101/2020.10.28.359927

**Authors:** Francois F. Barbier, Da Cao, Franziska Fichtner, Christoph Weiste, Maria-Dolores Perez-Garcia, Mathieu Caradeuc, José Le gourrierec, Soulaiman Sakr, Christine A. Beveridge

**Author notes:** **CORRESPONDING AUTHORS:** Francois F. Barbier, Christine A. Beveridge.

## Abstract

- Plant architecture is controlled by several endogenous signals including hormones and sugars. However, only little is known about the nature and roles of the sugar signalling pathways in this process. Here we test whether the sugar pathway mediated by HEXOKINASE1 (HXK1) is involved in the control of shoot branching.
- To test the involvement of HXK1 in the control of shoot architecture we modulated the HXK1 pathway using physiological and genetic approaches in diverse plants, rose, arabidopsis and pea and evaluated impacts of hormonal pathways.
- We show that triggering a hexokinase-dependent pathway was able to promote bud outgrowth in pea and rose. In arabidopsis, both HXK1 deficiency and defoliation led to decreased shoot branching and conferred hypersensitivity to auxin. *HXK1* expression was positively correlated with sugar availability. HXK1-deficient plants displayed decreased cytokinin levels and increased expression of *MAX2* which is required for strigolactone signalling. The branching phenotype of HXK1-deficient plants could be partly restored by cytokinin treatment and strigolactone deficiency could override the negative impact of *HXK1* deficiency on shoot branching.
- Our observations demonstrate that a HXK1-dependent pathway contributes to the regulation of shoot branching and interact with hormones to modulate plant architecture.

## INTRODUCTION

Crop yields are tightly correlated with the degree of shoot branching (Guo *et al.*, 2020), a trait which is mainly determined by the activation of formerly dormant axillary buds to produce branches (Evers *et al.*, 2011; Wang *et al.*, 2018). The high plasticity of this process allows plants to optimize their architecture to a given environment (Leduc *et al.*, 2014; Rameau *et al.*, 2015; Schneider *et al.*, 2019; Barbier *et al.*, 2019b). In order to acheive this plasticity, bud outgrowth is regulated by a complex endogenous signalling network involving hormones and nutrients including sugars (Domagalska & Leyser, 2011; Rameau *et al.*, 2015; Barbier *et al.*, 2019b).

The term ‘apical dominance’ is used to describe the growth control of the fast-developing shoot tip over axillary buds (Sachs & Thimann, 1964; Barbier *et al.*, 2017). Decapitation of the shoot apex leads to the outgrowth of the axillary buds at lower nodes. This phenomenon is mediated by two main mechanisms.

First, the very young leaves growing at the shoot tip produce a flow of auxin transported basipetally along the main stem to the root (Rameau *et al.*, 2015; Barbier *et al.*, 2017; Wang *et al.*, 2018). Auxin flowing in the stem inhibits bud outgrowth in an indirect manner since it cannot move into the bud (Hall & Hillman, 1975; Prasad *et al.*, 1993). Auxin indirectly acts on buds by reducing the levels of cytokinins (CK), which induce bud outgrowth, and presumably by promoting the synthesis of strigolactones (SL) that inhibit bud outgrowth (Brewer *et al.*, 2009; Hayward *et al.*, 2009; Domagalska & Leyser, 2011). The SL signal is perceived and transduced by a SCF complex involving the F-box MORE AXILLARY GROWTH2 (MAX2) and the α/β-hydrolase DWARF14 (D14). Upon SL binding to D14, MAX2 recruits an E3 ubiquitin ligase leading to the degradation of the uncharacterised SMXL6/7/8 proteins, which promote shoot branching (Chevalier *et al.*, 2014; Soundappan *et al.*, 2015; Waters *et al.*, 2017).

Second, the strong demand for nutrients by the growing shoot tip deprives the plant of the mobile sugars that are required to trigger bud outgrowth in pea and rose (Mason *et al.*, 2014; Barbier *et al.*, 2015a; Fichtner *et al.*, 2017). Notably, increased sugar availability alleviates the negative effect of auxin and strigolactone on bud release in pea, rose and chrysanthemum (Dierck *et al.*, 2016; Bertheloot *et al.*, 2020) indicating that sugars and auxin are involved in modulating bud outgrowth through an antagonistic action (Bertheloot *et al.*, 2020). Altering sugar partitioning by genetic manipulation of sugar transporters has also recently been reported to modulate shoot architecture in arabidopsis (Patzke *et al.*, 2019; Otori *et al.*, 2019). While the hormonal signalling pathways controlling shoot branching have been intensively studied over the past decades, the role of sugars in in process has only recently gained substantial attention (Van den Ende, 2014; Evers, 2015; Barbier *et al.*, 2015a). Besides their importance as a source of energy and carbon, sugars also function as signalling molecules to regulate plant growth and development (Rolland *et al.*, 2006; Lastdrager *et al.*, 2014; Sakr *et al.*, 2018). Sucrose is the main long-distance transported sugar in angiosperms (Lemoine *et al.*, 2013). It is proposed that sucrose has a signalling function in bud outgrowth (Mason *et al.*, 2014; Barbier *et al.*, 2015b,a; Fichtner *et al.*, 2017). In this respect, the low abundance disaccharide, trehalose 6-phosphate (Tre6P), which specifically reflects sucrose levels in plants and has a signalling function (Fichtner *et al.*, 2020; Fichtner & Lunn, 2021), was recently found to rapidly accumulate in pea bud outgrowth in response to decapitation (Fichtner et al., 2017), and to trigger shoot branching in arabidopsis (Fichtner *et al.*, In press), providing mechanistic evidence of a sucrose-specific signalling cascade in bud release.

Glucose and fructose, the sucrose cleavage products, also play signalling roles in plants. These two hexoses were also reported to trigger bud outgrowth (Rabot *et al.*, 2012; Salam *et al.*, 2017). However, their involvement in a sugar-signalling pathway has not been reported yet for bud outgrowth. Like sucrose, glucose also plays a signalling role in plant development (Li & Sheen, 2016) partly through HEXOKINASE1 (HXK1), the first enzyme of glycolysis, which converts glucose into glucose 6-phosphate. Besides its catalytic activity, the conformational change of HXK1 upon glucose binding generates a signal (Feng *et al.*, 2015; Wang *et al.*, 2017) that affects plant development independently of glucose phosphorylation (Moore *et al.*, 2003). The signalling activity of HXK1 is at least partly due to its ability to regulate gene transcription through binding DNA with other partners (Cho *et al.*, 2006). It also occurs through stabilizing certain transcription factors (Hu *et al.*, 2016).

Genetic evidence also supports a signalling role of HXK1. Mutation of *HXK1* leads to glucose insensitivity that can be recovered by complementation with a mutant version of HXK1 lacking catalytic activity (Moore *et al.*, 2003). This signalling function is not limited to HXK1 but also applies to other HXKs and HXK-like proteins lacking catalytic activity (Karve *et al.*, 2008, 2010; Karve & Moore, 2009).

Additional studies using glucose analogues support a signalling role for the HXK pathway. For example, mannose, a glucose analogue with a strong affinity for HXK and which is very slowly metabolised (Herold & Lewis, 1977), triggers the same response as glucose in a range of developmental processes (Chen & Jones, 2004; Hei *et al.*, 2018). On the other hand, 3-O-methylglucose (3-OMG), a slowly metabolised glucose analogue that can enter the cell but which has a poor affinity for hexokinase (Cortès *et al.*, 2003), is often used to highlight HXK-independent effects of glucose (Chen & Jones, 2004; Hei *et al.*, 2018).

Despite the fact that different soluble sugars can trigger shoot branching, only sucrose has been reported to play a signalling role in this process, being, at least partly, mediated by the Tre6P-dependent pathway. In this study we evaluate whether the HXK1-dependent pathway, specific to hexoses, contributes to the regulation of shoot branching. To achieve this, we used a combination of pharmacological and genetic approaches in different perennial (rose) and annual eudicot plants (pea and arabidopsis), in which a role of sugars in the control of shoot branching has been shown.

## MATERIAL AND METHODS

### Plant growth conditions

Rose plants were propagated by cutting and cultivated in a glasshouse. Pea plants were grown from seeds in a growth chamber (16hrs light, 20/22°C day/night). Arabidopsis seeds were first stratified for 72hrs at 4°C and transferred to a growth chamber with the same growth conditions as pea plants. In some experiments, arabidopsis plants were subjected to a 16hr night extension (light period replaced by a dark period) or to low (70±10 μmol m^−2^ s^−1^) and high light intensity (150±20 μmol m^−2^ s^−1^).

The *gin2-1* and *hxk1-3* knockout mutants in the arabidopsis Landsberg *erecta* (Ler) and Columbia-0 (Col-0) backgrounds, respectively, were obtained from NASC (N6383 and N69135) and have been previously described (Moore *et al.*, 2003; Huang *et al.*, 2015). The mutants *max4-10* and *max2-8* in Ler background were previously described (Nelson *et al.*, 2011). The homozygous double mutants *gin2-1/max4-10* and *gin2-1/max2-8* were selected in the F2 progeny based on the bushy phenotype of the plants for the *max4* and *max2* mutations, and trough sequencing of the *gin2* locus with the primers given in Supp. Table 1. The *35S::HXK1-31* line was provided by Prof. David Granot and has been described previously (Kelly *et al.*, 2012). The *35S::HXK1-32* line which was reported to have a bushy phenotype (Kelly *et al.*, 2012; Barbier *et al.*, 2015a) was not included this study after identifying a secondary mutation on *MAX4* coding sequence (data not shown).

### Shoot branching phenotyping in arabidopsis

For this study, we monitored the outgrowth of arabidopsis rosette axillary buds at an early and a flowering stage. For the early stage, the longest leaf primordia of buds were measured on plants 17-25 days after sowing. At this early stage, buds were usually 0.05-3 mm long and were measured under a dissecting microscope using a graticule eyepiece buds (Supp. Fig. S1A-B). The flowering stage was defined here by the elongation of the floral stem emerging from the axillary rosette buds, usually few days after the bolting of the main rosette (Supp. Fig. S1C). The branching at the flowering stage was phenotyped by scoring the number of rosette axillary branches with a bolt longer than 5 mm (Supp. Fig. S1C). Time-course of branch emergence was scored from the bolting date to prevent any effect due to development delay between genotypes.

### Decapitation experiments

Decapitation, defoliation and sugar supply in rose were performed by removing the shoot part above the fourth node with a true expanded leave on plants with a floral visible bud emerging from the apex, by removing the remaining leaves, and by supplying sugars in solution to the decapitated stump. Decapitation experiments in arabidopsis were performed by cutting the floral stem 5cm above the rosette and by the cauline and growing rosette buds (≥5mm) of plants with a 5-10 cm bolt. NAA (1-naphthaleneacetic acid, a synthetic auxin analogue) was applied in lanolin to the decapitated stump at a concentration of 5000 ppm (~ 27 mM). Lanolin without auxin, but supplemented with solvent, was applied to the stump of decapitated control plants. The decapitated stumps were trimmed every 5 days and the lanolin mix (with and without NAA) was refreshed to ensure correct auxin flow.

### *In vitro* split plate assay

The *in vitro* split plate assay was performed as described in Barbier *et al*. (2015). For the three species, excised stem segments bearing one node with a dormant axillary bud were briefly disinfected in 70% ethanol and cut to a length of 15 mm with the bud in the centre. Explants were then placed on a 1% agar medium supplemented with full strength MS (Duchefa) and with different sugar and auxin concentrations as described. Images were taken daily and bud length was measured using ImageJ (Schneider *et al.*, 2012). For pea, nodes were excised at the 5^th^ or 6^th^ nodal position on plants with 5 fully expanded leaves. For rose, nodes were excised from the second node bearing a true leaf of plants where the floral bud was just emerging from the shoot tip. For arabidopsis, nodes bearing a bud smaller than 1mm were collected on rosette and cauline branches.

### Cytokinin treatment

Cytokinin treatments were performed on 20-day old arabidopsis plants. The core of the rosette was treated with 15μl BAP (benzylaminopurine, a synthetic cytokinin analogue) dissolved in 50% ethanol (v/v) with 0.02% Tween-20 every three days until the plant started to senesce. The volume of BAP was progressively increased to 50μl as the plants grew. Control plants were treated with a mock solution (50% ethanol (v/v), 0.02% Tween-20).

### Gene expression analysis

For gene expression analysis in arabidopsis, the core of the rosette was harvested three weeks after sowing, frozen in liquid nitrogen and ground to a fine powder with an automated tissue grinder (Geno/Grinder^®^, SPEX). Total RNA was extracted as described in Barbier *et al*., 2019a using a simplified CTAB/PVP based method without phenol or chloroform. The diluted cDNA was used as a template for quantitative Real-Time PCR following the manufacturer’s instructions (SYBR Green, BIO-RAD) and product amplification was monitored with a CFX384 Touch™ Real-Time PCR Detection System (Bio-Rad). *TUBULIN3* and *Beta-ACTIN* were used for normalization. Primers used in this study are listed in Supp. Table 1.

### Plant hormone profiling

A part of the ground rosette material used for gene expression analysis was also used to extract hormones for quantification in arabidopsis. The extraction method was adapted from Cao et al. (2016). The samples were extracted with 80% acetonitrile, with 1% acetic acid containing d_5_-indole-3-acetic acid, d_6_-abscisic acid, d_5_-*trans*-zeatin, d_2_-dihydrozeatin, d_5_-*trans*-zeatin-riboside, d_3_-dihydrozeatin-riboside, d_5_-zeatin-riboside-5-monophosphate, d_6_-isopentenyladenine, d_6_-isopentenyladenosine, d_6_-isopentenyladenine-7-glucoside, and d_6_-isopentenyl-adenosine-5-monophosphate as internal standards (n.b. 9 Zeatin-9-glucoside and isopentenyladenine-7-glucoside were quantified relative to the wild-type). Samples were purified with Sep-Pak tC18 cartridges (Waters Corporation) and then used for LC-MS measurements. The LC-MS system was a Nexera X2 liquid chromatograph (UHPLC) system (Shimadzu Corporation) coupled with a 5500 QTRAP MS system (AB Sciex). The extracted hormones were separated on a Phenomenex Kinetex C18 reversed phase column (2.1mm × 100 mm, 1.7 μm).

### Statistical analyses

Significant differences between samples were determined using a Student’s *t*-test or with a one-way ANOVA (analysis of variance) when appropriate (with α=0.95 / *p*-value <0.05). Significant differences are represented by asterisks (Student’s *t*-test) or letters (one-way ANOVA), respectively. For interaction tests in Supplementary Table 1, a two-way ANOVA was used. In all figures, data is given as means +/− standard error (se).

## RESULTS

### HXK-phosphorylated hexoses trigger bud outgrowth *in vitro*

We previously reported that feeding sucrose to decapitated and defoliated rose plants promotes axillary bud outgrowth (Bertheloot *et al.*, 2020). In order to assess whether a HXK-dependent pathway could be involved in sugar-induced bud outgrowth, we fed decapitated and defoliated plants with sucrose, the main long-distance transported sugar in most plants, and mannoheptulose (MHPT), a competitive inhibitor of HXK (De la Fuente *et al.*, 1958; Coore & Randle, 1964; Pego *et al.*, 1999) (Fig. 1A). We observed an increase in bud size 7 days after the beginning of the treatment when plants were supplied with sucrose compared with intact plants and plants fed with mannitol as an osmotic control (Fig. 1B). In contrast, MHPT treatment significantly inhibited the sucrose-induced bud outgrowth (Fig. 1B), suggesting that a HXK-dependent pathway may be involved in the control of shoot branching by sugars.

**Figure 1.**
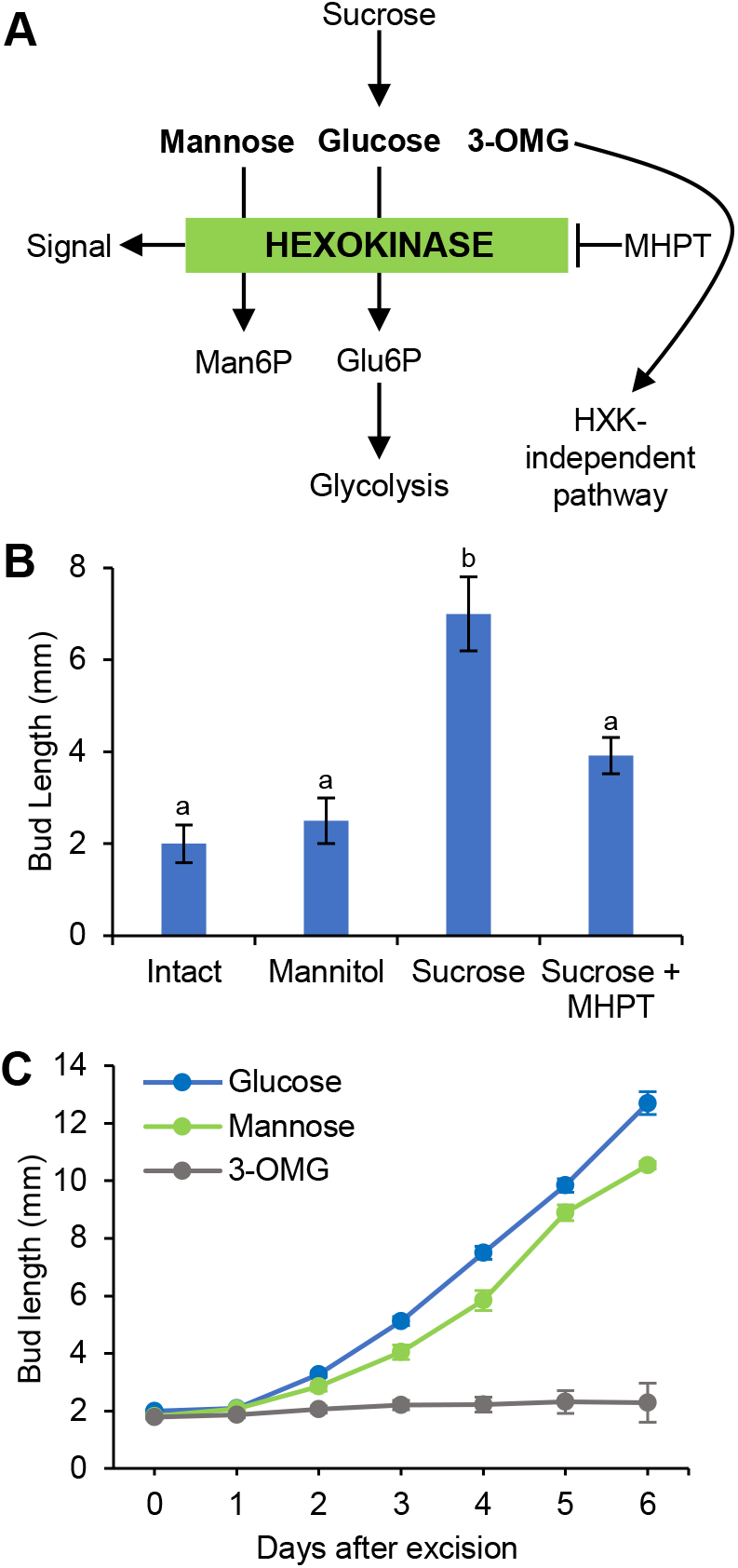
Impact of hexokinase inhibitor and glucose analogues on bud elongation in rose. **A**. Schematic representation of the impact of mannoheptulose (MHPT) on hexokinase and of the metabolism of glucose analogues (in bolt). **B**. Impact of 30 mM mannitol, 30 mM sucrose and 30 mM sucrose + 50 mM MHPT on bud elongation in decapitated and defoliated plants after 7 days of supply through the decapitated stump and compared to intact plants. Data are mean ±se of the bud below the decapitation site (n=6). Letters indicate significant differences between each condition (one way ANOVA). **C**. Impact of 30 mM glucose, mannose and 3 O-methylglucose (3-OMG) on single bud elongation *in vitro*. Data are mean ±se (n=8-10).

If the HXK-dependent pathway is important in bud outgrowth, we would predict that substrates of HXK would trigger bud outgrowth. We therefore supplied excised single nodes from rose and pea with different sugars *in vitro* (Fig. 1C, Supp. Fig. S2A). Glucose, the main substrate of HXK1 in plants, quickly triggered the buds to grow out in both species, while galactose, a hexose metabolised independently of the HXK-mediated step, could not trigger the buds to grow out after 6 days of incubation (Fig. 1C, Supp. Fig. S2A). Two slowly-metabolisable glucose analogues, mannose and 3 O-methylglucose (3-OMG), also produced responses expected for a role of HXK1 in bud outgrowth. These two sugars both enter the cell, but while mannose is preferentially phosphorylated by HXK, 3-OMG has poor affinity with HXK (Fig. 1A). Like glucose, mannose could induce bud growth, whereas no elongation was observed with buds fed with 3-OMG (Fig. 1C). The same result was observed with pea (Supp. Fig. S2A), suggesting that this effect is not specific to rose, and is conserved between these perennial and annual plants.

To provide more evidence that mannose acts independently of its metabolism for growth, we measured the dry weight of single nodes (stem and buds together) 3 days after glucose, mannose and 3-OMG feeding (Supp. Fig. S3A). Under these conditions, glucose and, to a lesser extent, 3-OMG increased the dry matter of the single nodes, whereas mannose did not, suggesting that mannose was not used as a carbon source in this system. The fact that mannose increased bud growth but no change in dry weight, whereas 3-OMG did not increase bud growth but increased dry weight (Fig. 1C and S3A), further suggests that sugars are required as a signal and not necessary as a carbon or energy source to promote bud release. Additionally, the effect of mannose on bud outgrowth was independent of its concentration (Supp. Fig. S3B), which is inconsistent with an involvement in a metabolic pathway where a dose response would be expected, as has previously been reported with metabolised sugars used in the same range of concentrations (Rabot *et al.*, 2012; Mason *et al.*, 2014; Barbier *et al.*, 2015b; Fichtner *et al.*, 2017). To exclude the possibility that the absence of bud outgrowth in presence of 3-OMG was due to a potential toxic effect of this analogue, we cultivated single nodes in a combination of sucrose and 3-OMG. The result indicated that 3-OMG did not inhibit the sucrose-induced bud outgrowth (Supp. Fig. 3C), suggesting that this analogue is not toxic for rose buds in the concentration used in our assays. Altogether, these results suggest that the regulation of bud outgrowth involves a HXK-dependent pathway that is largely independent of HXK1 metabolic activity.

### Change in sugar availability regulates shoot branching and *HXK1* expression in arabidopsis

The results obtained in rose and pea suggested that a HXK-dependent pathway may hold a signalling function in the control of vegetative bud outgrowth (Fig. 1 and Supp. Fig. S2). As HXK1 from *Arabidopsis thaliana* was reported to play a role in sugar signalling (Moore *et al.*, 2003), we investigated whether HXK1 could be involved in shoot branching and bud outgrowth in this species. Although shoot branching in arabidopsis has been shown to be regulated by sugar availability at a flowering stage (Barbier *et al.*, 2015b; Otori *et al.*, 2017, 2019; Patzke *et al.*, 2019), no studies have tested this at an early stage. Since vegetative and floral buds may have different environmental and genetic cues regulating their outgrowth, it is important to test the response of buds to sugar availability at an early stage in arabidopsis in order to compare to the results obtained in rose and pea. We therefore monitored the elongation of rosette axillary buds at an early stage, when axillary buds were really young and only leaf primordia were visible. We modulated endogenous sugar levels through differential light intensity and night extension (Smith & Stitt, 2007; Usadel *et al.*, 2008; Moraes *et al.*, 2019) and measured the length of axillary buds at the early stage in young rosettes (Fig. 2A-B and Supp. Fig. S1A-B). Our results show that one-night extension and low light intensity both led to a reduced elongation of all axillary buds along the main axis of the plant (Fig. 2A-B), supporting the contention that sugar availability modulates bud outgrowth at an early stage in arabidopsis.

**Figure 2.**
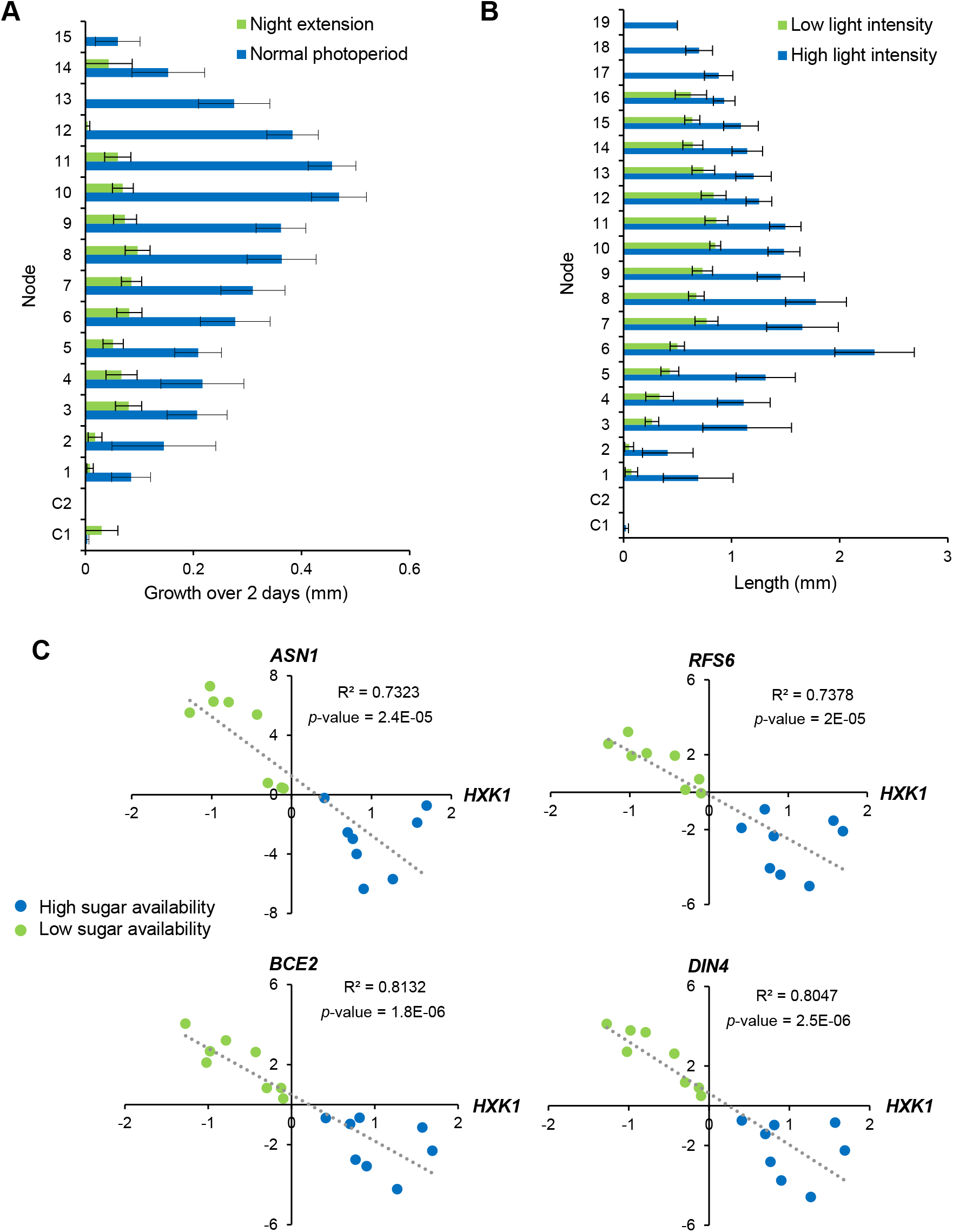
Impact of sugar availability on shoot branching and *HXK1* expression in arabidopsis. **A**. Plants exposed to a one-night extension or to regular photoperiod at day 23 after sowing. Bud elongation measured over following 2 days. **B**. Plants grown continually under low light (70±10 μmol m^−2^ s^−1^) or high light intensity (150±20 μmol m^−2^ s^−1^). Buds were measured when the rosette diameter was similar under the two conditions (low light, 32 d; high light, 27 d). For **A** and **B** data are mean ±se (n=12-15) for buds at different nodal positions numbered acropetally from the cotyledonary node (marked C1, C2) and grown under 16h light and 8h night. **C**. Correlation between *HXK1* expression and sugar marker genes (*ASN1* (*DIN6*, AT3G47340), *RS6* (*DIN10*, AT5G20250), *BCE2* (*DIN3*, AT3G06850) and *DIN4* (AT3G13450)) in different published transcriptomic analyses from experiments with varied endogenous sugar levels (Supp. Fig. S5). Values displayed are Log2. Low and high sugar availabilities (green and blue circles, respectively) have been determined by the expression of the sugar marker genes (positive value = low sugar availability, negative value = high sugar availability).

After showing that sugar availability regulates early bud outgrowth in arabidopsis, we used publically available transcriptomic data sets (Genevestigator; Hruz et al., 2008) to explore the response of *HXK1* expression to diverse signals controlling shoot branching. We first selected treatments leading to elevated or reduced sugar levels in plants, including the conditions mentioned above. We then correlated *HXK1* expression with the expression of four broadly used starvation markers (Fig. 2C and Supp. Fig. S4A). The results indicate a strong and significant negative correlation between the expression of *HXK1* and starvation marker genes (Fig. 2C). These observations indicate that *HXK1* expression is positively regulated by sugar availability in arabidopsis.

In order to test whether HXK1 could act downstream of hormones involved in shoot branching we determined the response of *HXK1* expression to diverse hormones. The dataset displayed in Supp. Fig. S4B highlights the fact that *HXK1* expression in seedlings is not responsive to hormone treatments including auxin, cytokinins, abscissic acid, gibberellins, ethylene or brassinosteroids. Moreover, *HXK1* expression was only significantly regulated by two perturbations among the 135 perturbations corresponding to the available set of hormone treatments in Genevestigator (Supp. Fig. S4C). Altogether, these results suggest that *HXK1* expression is largely not responsive to hormones and is instead specifically regulated by sugar availability.

### HXK1 modulates shoot branching in arabidopsis

The results obtained in the previous experiments suggest that HXK1 may be involved in the regulation of shoot branching in arabidopsis. We tested this at the genetic level by characterizing the branching phenotype of the *glucose insensitive 2-1* (*gin2-1*) mutant, an arabidopsis *hxk1* knockout mutant (Moore *et al.*, 2003). We measured the elongation of the axillary buds at an early stage on different days prior to rapid elongation of the primary floral stem (bolting) (Supp. Fig. 1A-B). We then characterized shoot branching in arabidopsis when rosette axillary buds were at the flowering stage by scoring the number of primary rosette axillary inflorescences longer than 5mm on different days after bolting (Supp. Fig. 1C). Compared to its wild-type control (Landsberg *erecta*, Ler), the growth of the *gin2* buds at the fourth nodal position was strongly delayed at an early stage (Fig. 3A). This inhibition was also observed at different nodal positions (data not shown). Since *gin2* displays a smaller rosette diameter than L*er* at this early stage, we normalised the bud length to rosette diameter to avoid any bias due to a potential difference in the photosynthetic leaf area. The result demonstrated that *gin2* still displayed an inhibition of bud elongation at the early stage (Supp. Fig. S5). *Gin2* mutant plants also had delayed timing of branch elongation and decreased final branch number (~50% less compared to Ler, Fig. 3B-C). Similar to *gin2* plants in the L*er* background, *hxk1* plants in Col-0 had significantly fewer rosette axillary branches than wild-type plants (Fig. 3D). This genetic evidence therefore shows that HXK1-deficiency negatively impacts shoot branching in arabidopsis at broad stages of plant ontogeny.

**Figure 3.**
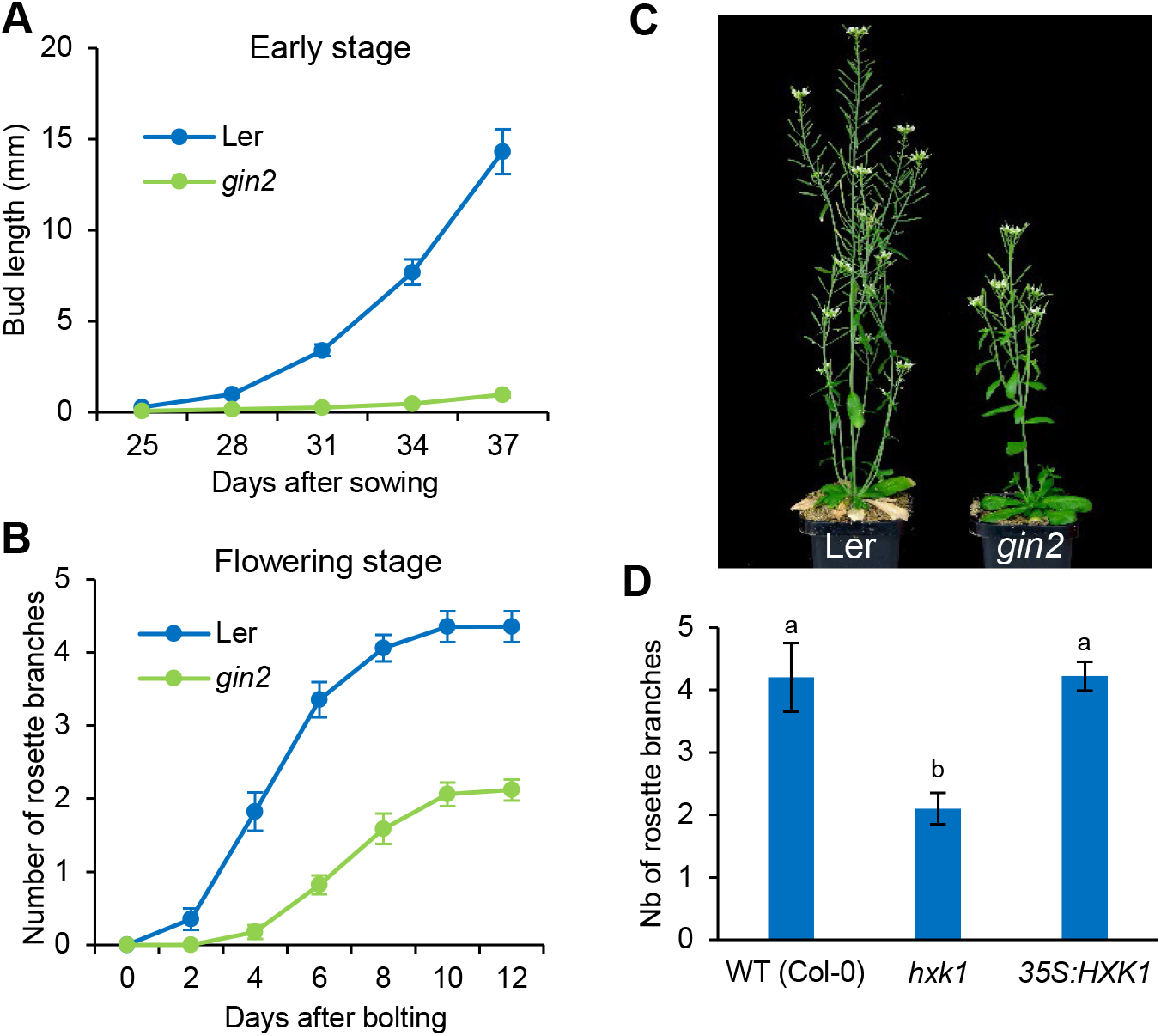
Impact of HXK1 deficiency on shoot branching in arabidopsis. **A** Length of the node 4 bud at the early stage and **B.**, number of rosette branches ≥ 5 mm at the flowering stage in *gin2-1* and wild-type (Ler); Data are mean ±se (n=15). **C**. Ler and *gin2-1* phenotype at flowering stage. **D** Final number of rosette branches at the flowering stage in wild-type (WT; Col-0), *hxk1-3* mutant and HXK1 over-expresser (*35S:HXK1*). Data are mean ±se (n=12-15). Letters indicate the significant differences between the genotype determined with a one-way ANOVA.

In order to test whether excessive HXK1 activity may cause the converse, an increase in branching, we phenotyped plants over-expressing HXK1 (*35S:HXK1-31*). In this case, there was no significant effect of HXK1 over-expression on the number of rosette axillary branches compared with wild-type controls (Fig. 3D). This indicates that HXK1 is required to maintain branching to the wild-type level in intact plants and that overexpression of HXK1 cannot over-ride the effect of apical dominance.

### HXK1 deficiency enhances auxin response

Apical dominance studies involving shoot tip removal have been used widely to test the role of auxin in shoot branching and more recently in demonstrating a role of sugars (Sachs & Thimann, 1964; Cline, 1996; Morris *et al.*, 2005; Mason *et al.*, 2014; Barbier *et al.*, 2019b; Bertheloot *et al.*, 2020). In order to get insight into the role of HXK1 in apical dominance in arabidopsis, we compared the branching phenotype of L*er* and *gin2* in response to different treatments that modulate auxin and/or sugar levels: decapitation, auxin application and defoliation (Fig. 4). It took six days longer for the *gin2* mutant to initiate the first branches compared to L*er*, and it took longer for the decapitated *gin2* plants to reach the plateauing phase compared with the wild-type (Fig. 4A-B). To highlight the differences between the two genotypes, we compared the number of branches at the beginning of the plateauing phase (12d for Ler and 24d for *gin2*, Fig. 4A-C). As observed in Figure 3, intact *gin2* plants displayed a decreased branching phenotype compared to the wild-type control. Decapitation led to an increased branching phenotype in both genotypes but *gin2* plants still displayed a significantly lower number of branches than Ler in response to this treatment (Fig. 4C). However, compared with wild-type the inhibitory effect of *gin2* on branching was reduced in decapitated plants (25% inhibition) compared with intact plants (50% inhibition). Moreover, the enhancement of branching by decapitation was higher in *gin2* (440% increase) than in Ler (310% increase) (Fig. 4C). These results show that the effect of *gin2* on shoot branching is less pronounced in decapitated plants than in intact plants.

**Figure 4.**
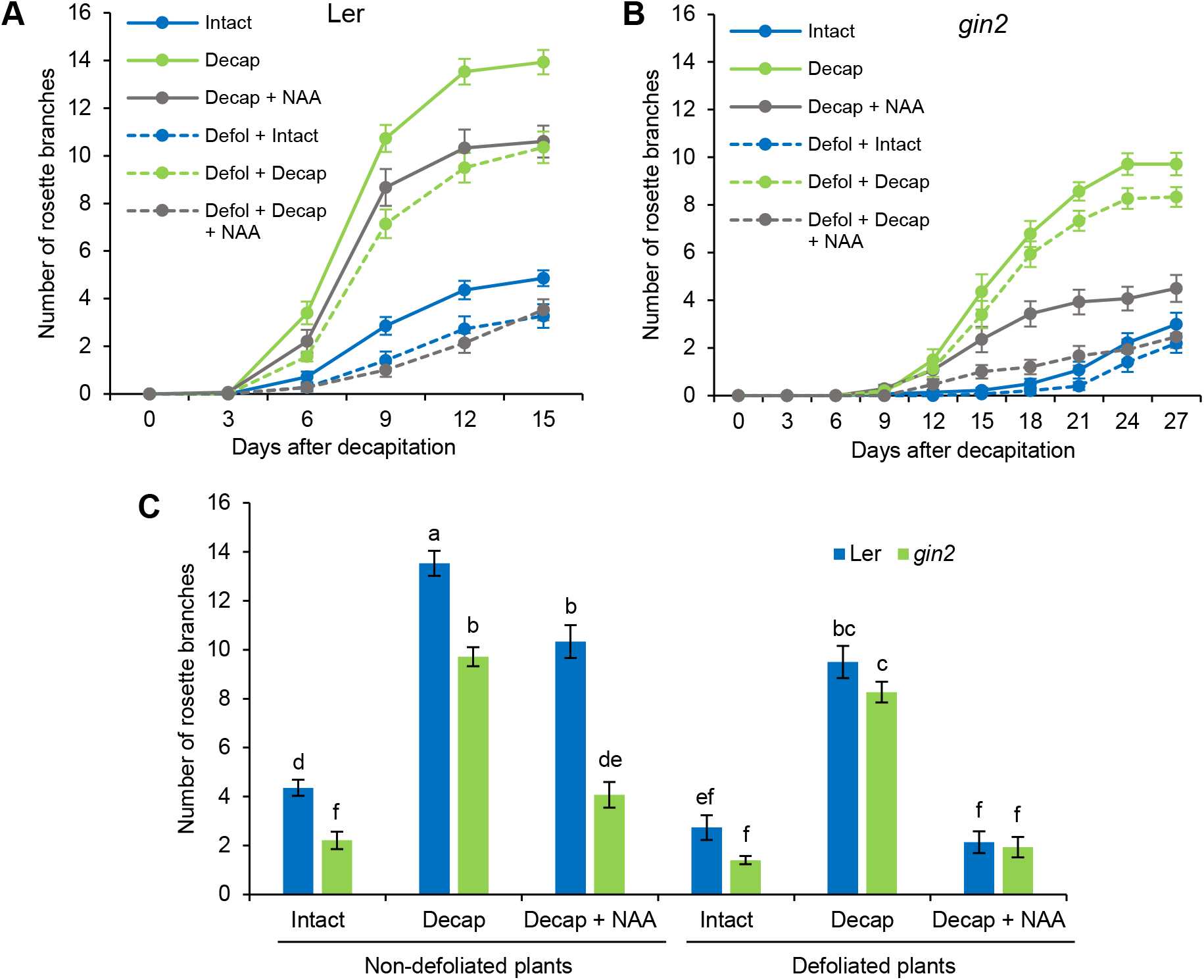
Impact of HXK1 deficiency on apical dominance and decapitation response in *Arabidopsis*. Number of rosette branches ≥ 5 mm in **A.**, *wild-type* (Ler) and **B.**, *gin2-1* in controls (intact), and decapitated plants ± 5000 ppm applied to the decapitated stump and ± defoliation and grown under the same conditions. Data are mean ±se (n=12-15). **C**. Number of rosette branches ≥ 5 mm when branch emergence started to plateau (12 and 24 days after bolting for Ler and *gin2-1*, respectively).

Unlike what was previously reported in the Col-0, a high supply of auxin to the decapitated stump resulted in a partial restoration of apical dominance in non-defoliated Ler plants. In contrast, auxin treatment almost fully restored apical dominance in defoliated Ler plants (Fig. 4C). Interestingly, this trend was also observed in non-defoliated *gin2* plants. The ability of sugar or HXK1 deficiency to enhance auxin responsiveness supports the idea that auxin and sugar availability have an antagonistic effect on shoot branching in arabidopsis. In this way, HXK1 deficiency can be thought to mimic the effect of decreased sugar levels during shoot branching in intact plants.

To further support the hypothesis that defoliation modulates shoot branching by depriving buds from sugars and not from any other leaf-derived signal, we grew single nodes bearing a dormant bud with or without the adjacent leaf on a sugar-free medium *in vitro* (Fig. 5A). Under these conditions, the wild-type buds grew only in presence of the leaf. The addition of sucrose (30mM) to the growth medium was sufficient to restore bud outgrowth, supporting the hypothesis that the inhibitory effect of defoliation is due to a decrease in sugar availability (Fig. 5A).

**Figure 5.**
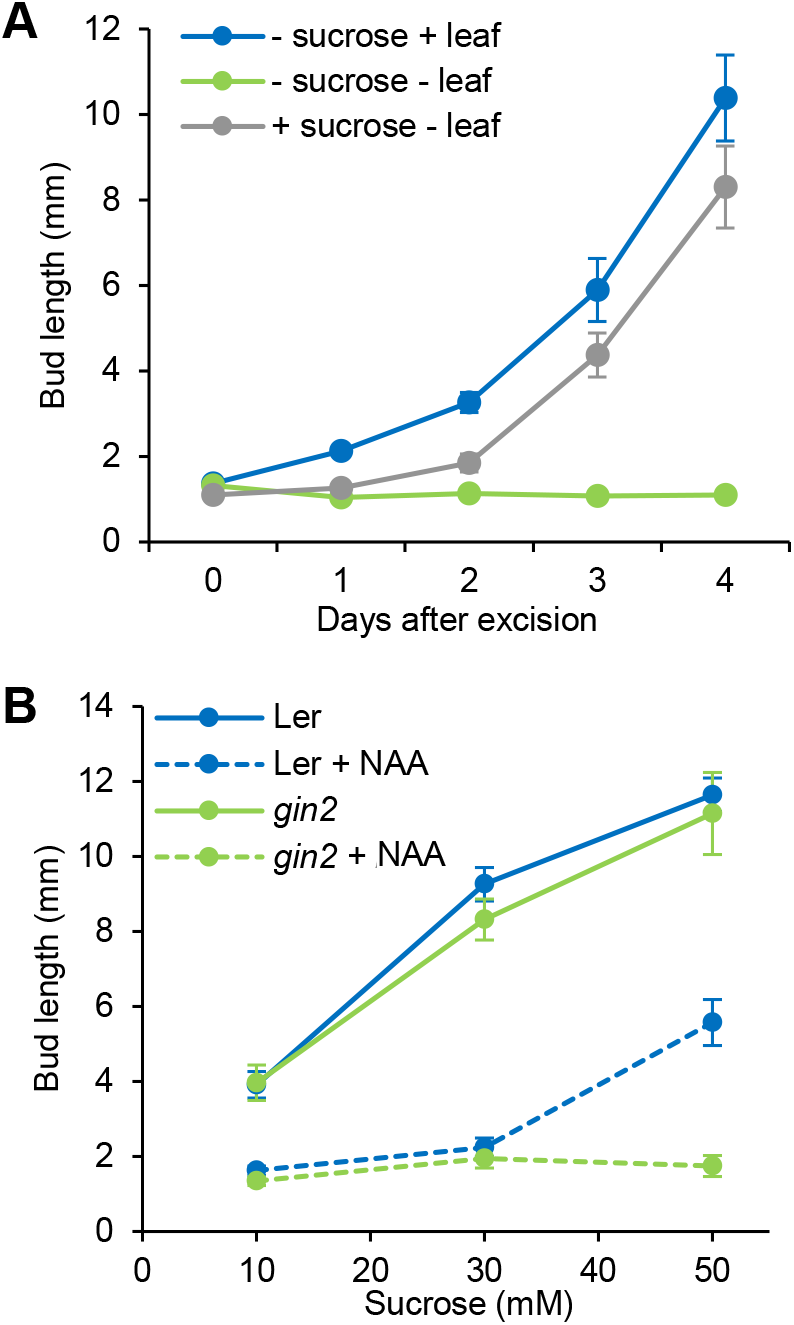
Impact of sugar availability and HXK1 deficiency on cauline buds *in vitro*. **A**. Bud growth of isolated wild-type (Col-0) cauline nodal stem segments with and without their adjacent leaf and ± 30 mM sucrose. Data are mean ±se (n=7-8). **B**. Length of wild-type (Ler) and *gin2-1* isolated cauline buds ± 2.5 μM NAA and 10, 30 or 50 mM sucrose, data are mean ±se (n=10-11) at d 6. Two-way ANOVA test is presented in Supplementary Table 2.

To further assess the role of HXK1 in mediating the effect sugar availability during antagonism between sugars and auxin during bud outgrowth in arabidopsis, we supplied cultivated single nodes with different concentrations of auxin and sucrose (Fig. 5B). With 30 mM sucrose, auxin could strongly inhibit bud elongation in the wild-type (76%). Increasing the sucrose concentration to 50 mM in the growth medium reduced this inhibitory effect of auxin (52%). Unlike the wild-type, auxin completely inhibited bud outgrowth in the *gin2* mutant regardless of sucrose concentration. Statistically analysis of these results indicate that the combined effect of sucrose, auxin and *gin2* mutation on buds is not additive and instead results from an interactive effect between these components (Figure 5B; Supp. Table 2). This supports the hypothesis that HXK1 mediates the effect of sugar availability to dampen the inhibitory effect of auxin on bud outgrowth thereby conferring increased sensitivity to auxin in the *gin2* mutant.

### The *gin2* mutant is cytokinin deficient

The results presented in Figures 3-5 highlight that HXK1 deficiency negatively impacts shoot branching in intact arabidopsis plants but has only a moderate effect in decapitated plants. This differing response of *gin2* mutant and WT under intact and decapitated conditions is reminiscent of those reported for arabidopsis CK-deficient mutants, which display a decreased branching phenotype in intact plants but not in decapitated plants (Müller *et al.*, 2015). We evaluated whether the HXK1-deficient mutant *gin2* is affected in the CK pathway by first measuring CK levels in the rosette of plants with a visible floral bud in the centre of the rosette. All of the *trans*-zeatin-type CKs including trans-zeatin riboside-5’-monophosphate, *trans*-zeatin riboside, *trans*-zeatin-9-glucoside, *trans*-zeatin as well as the isopentenyl-type CK, isopentenyladenosine-5’-monophosphate showed significantly lower content in *gin2* compared with L*er* (Fig. 6A). CK deficiency may therefore contribute to the phenotype of the *gin2* mutant.

**Figure 6.**
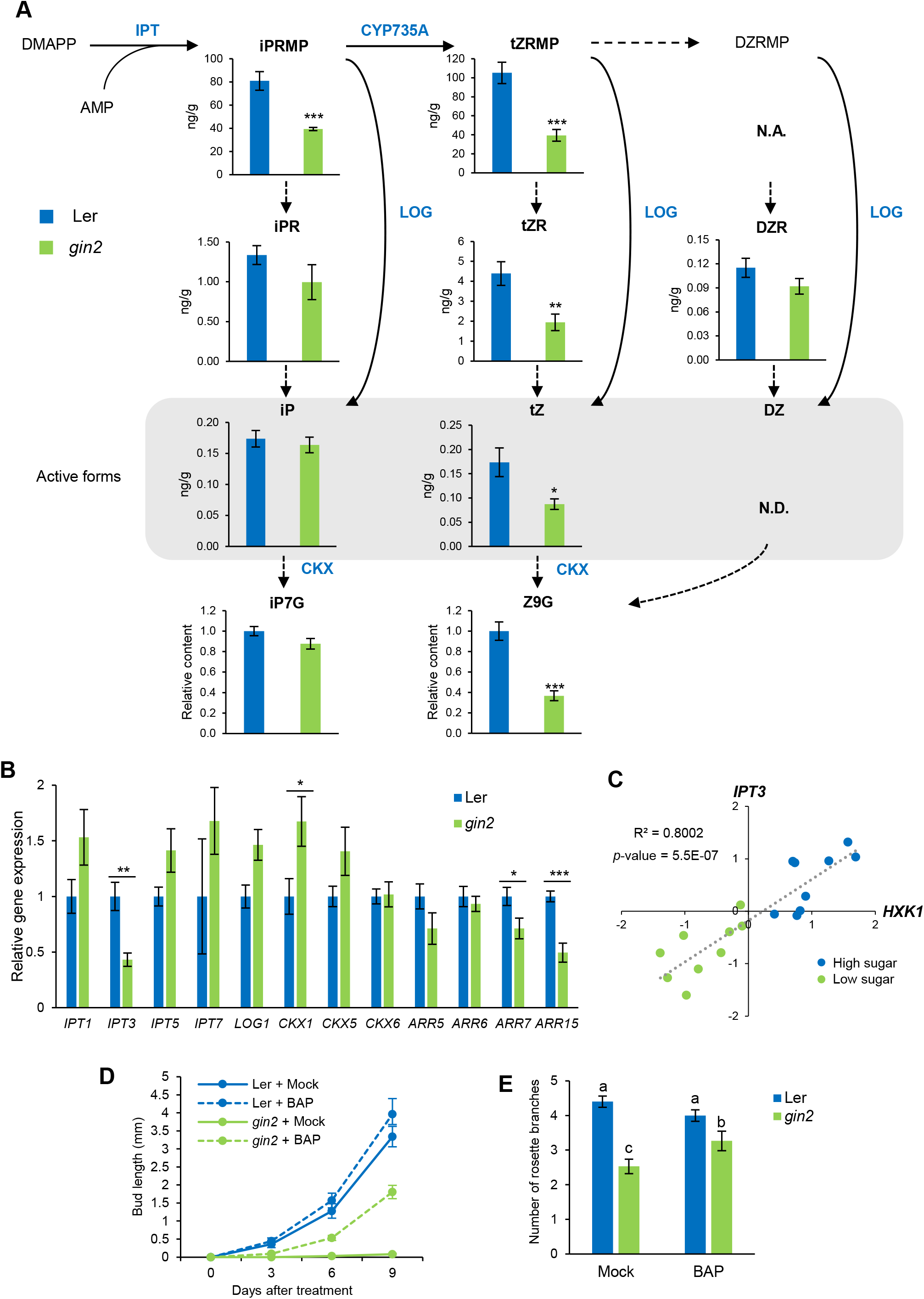
Impact of HXK deficiency of the cytokinin pathway. **A**, Cytokinin levels and **B,** cytokinin-related gene expression in wild-type (*Ler*) and *gin2-1*. Data are mean ±se (n=6 individual rosettes; *, *p*-value<0.05; independent t-test); whole rosettes were harvested when the first flower bud was visible (~4 weeks after sowing) **C**. Correlation between *HXK1* and *IPT3* expression in the same samples as in Figure 3C. Data displayed are Log2. **D**. Length of the node 4 rosette bud at the early stage and **E**. final number of rosette branches at the flowering stage ≥ 5 mm in Ler and *gin2-1* ± 50 μM BAP. Data are mean ±se (n=15). Letters indicate significant differences (one-way ANOVA).

The CK deficiency in *gin2* was reflected by differences in CK-related gene expression in the same tissue (Figure 6B). Among the tested CK biosynthesis- and degradation-related genes, *IPT3* expression was significantly decreased in *gin2* compared to the wild-type, while *CKX1* was significantly increased. To confirm that the decrease in CK levels has a negative impact on its downstream signalling, we monitored the expression of four *ARR* genes, which have been shown to be strongly responsive to CK in arabidopsis shoots (Potter *et al.*, 2018). As expected, the decrease of *ARR7* and *ARR15* expression in *gin2* reflected the decreased CK levels in this mutant. We also observed a positive correlation between sugar availability and the expression of *IPT3*, *ARR7* and *ARR15* observed across publically available transcriptomic data sets incorporating treatments that modulate endogenous sugar levels (Supp. Fig. S4A). Moreover, we observed a strong and significant positive correlation between the expression of *HXK1* with *IPT3* in response to these treatments (Fig. 6C).

To test whether the decreased branching phenotype of *gin2* could be due to a CK deficiency, we treated the axillary buds at the early stage (2-3 week old plants) with BAP (benzylaminopurine), a synthetic cytokinin analogue (Fig. 6D-E). BAP treatment did not increase bud elongation in Ler, but it significantly promoted bud elongation in *gin2* (Fig. 6D). Ongoing CK treatment slightly but significantly increased the final number of axillary branches scored in *gin2* but not in wild-type plants (Fig, 6E). Despite the increase in early axillary bud growth and final branch number, the BAP treatment to *gin2* did not restore branching to wild-type levels. Nevertheless, these observations demonstrate that the *gin2* mutant is lacking in bioactive CK, and that this deficiency is partly responsible for the decreased branching observed in this mutant.

### HXK1 action though the strigolactone pathway

We previously reported for rose and pea that sugars promote shoot branching by suppressing strigolactone perception (Bertheloot *et al.*, 2020). We also showed that the inhibitory effect of sugar depletion could be largely overcome by SL-depletion. We therefore tested whether impairing the SL pathway could overcome the inhibitory effect of *gin2* on shoot branching in arabidopsis. As predicted, the inhibition of shoot branching by *gin2* could be largely overcome by the inhibition of the SL pathway in both biosynthesis (*max4-1*) and signalling (*max2-1*) mutants (Fig. 7C). The inductive effect of *max4* and *max2* mutations on shoot branching was higher in the *max2* mutant than in the *max4* mutant (Fig. 7C). This difference between SL-deficient mutants and *max2* mutants has already been reported in different development and physiological processes (Ishikawa *et al.*, 2005; Umehara *et al.*, 2008; Hayward *et al.*, 2009; Rasmussen *et al.*, 2012; Chevalier *et al.*, 2014; Kalliola *et al.*, 2020), and it was shown that MAX2 retains a function on shoot branching independent of SL (Stirnberg *et al.*, 2007).

**Figure 7.**
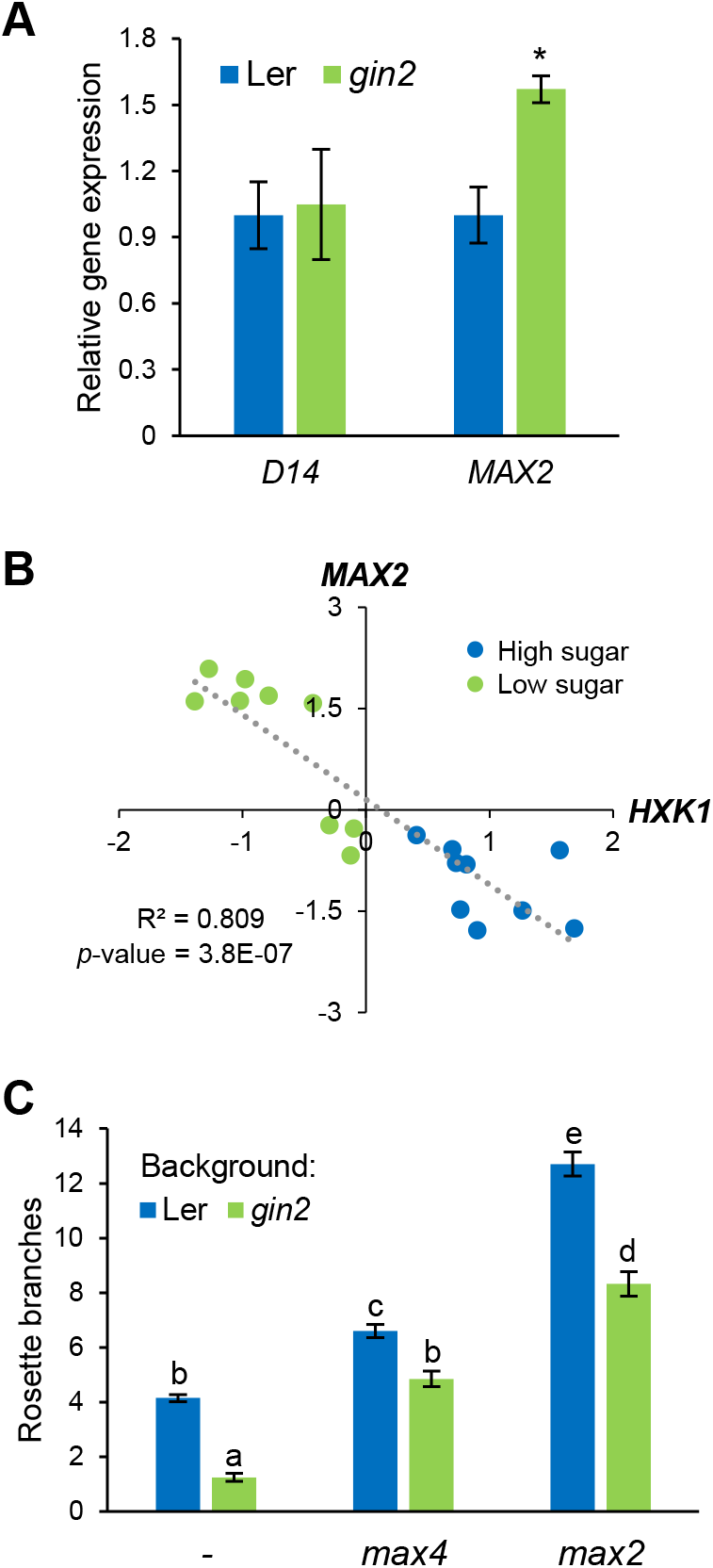
Impact of HXK1 and strigolactone deficiency on shoot branching in *Arabidopsis thaliana*. **A**. *D14* and *MAX2* expression in 4 week old rosettes. Data are mean ±se (n=6). Stars indicate significant differences between Ler and *gin2* (*, p-value<0.05; independent t-test) **B.** Correlation between *HXK1* and *MAX2* expression in the same samples as in Figure 3C. Data displayed are Log2. **C**. Final number of rosette branches longer than 5 mm in Ler, *gin2-1, max4-1, max2-1, gin2-1/max4-1 and gin2-1/max2-1*. Data are mean ±se (n=12-16). Letters indicate the significant different between the genotypes determined with a one-way ANOVA.

Based, on our previous work, we would predict that sugar availability and the HXK1 pathway may affect the expression of genes involved in SL perception and signalling. The present results show that *MAX2* expression is upregulated in the *gin2* mutant in arabidopsis (Fig. 7A). *MAX2* expression was also inhibited in pea buds fed with sucrose, glucose and mannose, but not 3-OMG feeding, (Supp. Fig. S2B). We also observed a negative correlation between sugar availability and the expression of *D14* and especially *MAX2* across transcriptomic data sets including treatments that modulate endogenous sugar levels (Supp. Fig. S4A). Additionally, *MAX2* expression also strongly negatively correlated with *HXK1* expression under these treatments (Fig. 7B). These results suggest that sugar availability and the HXK1-dependent pathway have a negative impact on the expression of *MAX2*, involved in strigolactone perception.

The results of this study provide evidence that the HXK1 pathway acts to promote branching via suppression of SL perception (Fig. 7) and induction of CK levels (Fig. 6). If this is indeed the case, we would then expect *gin2* to retain an inhibitory effect in the SL mutants because CK levels should be reduced by HXK1-deficiency. As expected, double mutants of *gin2* and strigolactone biosynthesis (*max4-1*) and signalling (*max2-1*) mutants both showed less branching than the strigolactone single mutants (Fig. 7C). This suggests that HXK1 does not solely act though SL.

Using a physiological approach to investigate the role of sugar availability and strigolactones on shoot branching, we observed only a one-night extension could also inhibit early bud outgrowth in the SL-deficient mutant *max4* (Supp. Fig. S6), but to a lesser extent than in the wild-type (Fig. 2A). Based on these findings, we suggest that inhibition of the SL pathway largely, but not totally, overrides the negative impact of sugar starvation and HXK1 deficiency on shoot branching.

## DISCUSSION

### HEXOKINASE1 mediates the sugar status of plants to modulate shoot architecture

Sugar availability is proposed as a critical factor in determining the shoot branching pattern in different species including arabidopsis (Evers, 2015; Otori *et al.*, 2017; Barbier *et al.*, 2019b). Here we show that the HXK1 pathway is required for shoot branching. HXK1 deficiency in arabidopsis causes decreased shoot branching. HXK is the first enzyme of the glycolysis, so the negative impact of the *hxk1/gin2* mutation could be attributed to a decreased carbon metabolism. The mannose-triggered bud outgrowth observed in rose and pea suggests that this HXK-dependent pathway may be involved in a signaling manner during shoot branching. Indeed, since mannose is not metabolized after its phosphorylation by HXK1, the effect of mannose on bud growth is likely to be due to HXK1 signalling. Further studies at the genetic level are required to fully demonstrate the signalling role of HXK1 in shoot branching.

Most studies on arabidopsis focus on the branching pattern at the flowering stage of axillary bud development. The present study demonstrates that sugar availability also modulates shoot architecture at an early stage, before the bolting of the axillary buds. Our results show that a HEXOKINASE-dependent pathway plays an important role in the control of shoot branching by sugar availability when buds are at an early stage of development, before the elongation of the main floral shoot, as well as when buds are at the flowering stage.

HXK1 has been reported to play a role in different developmental processes (Moore *et al.*, 2003; Granot *et al.*, 2014; Li *et al.*, 2018). For instance, HXK1 negatively regulates root meristem development in arabidopsis (Huang *et al.*, 2019), and has been found to have a positive role on cell proliferation and expansion during sucrose-induced leaf growth (Van Dingenen *et al.*, 2019). Similarly to arabidopisis, reduction of *HXK1* expression through silencing in tobacco leads to inhibition of leaf growth in source and sink tissues (Kim *et al.*, 2013). These observations are in accordance with our results showing the growth promoting role of HXK1 during shoot branching. In *Brassica napus*, *HXK1* expression in axillary buds has recently been shown to be positively correlated to the branching gradient along the axis (Li *et al.*, 2020). Our results also indicate that *HXK1* expression is inhibited by conditions leading to decreased sugar levels in arabidopsis (Fig. 2 and Supp. Fig. S4A). Regulation of *HXK1* gene expression may therefore play a significant role in mediating the sugar status of plants to modulate shoot branching, thereby providing means for plants to adjust their architecture to environmental conditions that affect the plant’s energy homeostasis (Fig. 8A).

**Figure 8.**
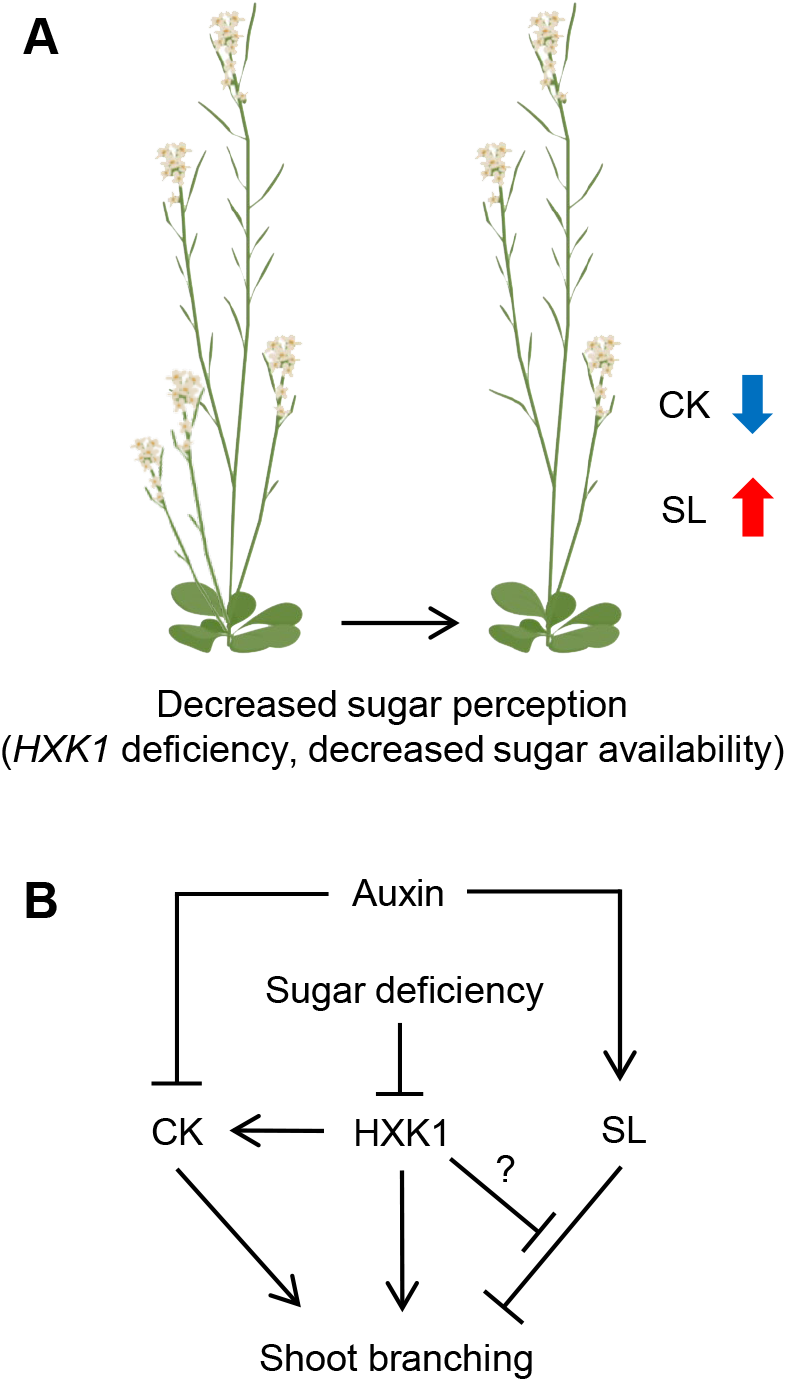
Proposed involvement of HXK1 during the control of shoot branching in intact plant. **A**. Decreased sugar levels and therefore reduced *HXK1* expression reduce shoot branching in intact plants. This is correlated with decreased CK levels and increased SL perception. Arrows indicate positive (blue) and negative (red) hormonal pathways involved in shoot branching. The direction of the arrows indicates up- and down-regulation. **B**. In intact plants, the low sugar availability for buds inhibits *HXK1* expression leading to decreased CK levels and potentially to an increased SL perception, thus increasing the inhibitory effect of auxin. A pathway independent of SL and CK cannot be excluded.

It was previously reported that a sucrose-specific pathway is potentially involved (Barbier *et al.*, 2015b; Salam *et al.*, 2017) and that Tre6P is a strong candidate for mediating this process (Fichtner *et al.*, 2017). Tre6P accumulates in pea axillary buds upon decapitation, and its increase correlates with bud elongation (Fichtner *et al.*, 2017). The present study demonstrates that a HXK1-dependent pathway, specific to hexoses, is also involved in the control of shoot branching. The role of the HXK1-dependent pathway in this shoot branching is likely different from the reported role of Tre6P in shoot branching. Indeed, contrary to HXK1 over-expression which had no measurable effect on shoot branching (Fig. 3), the over-expression of Tre6P-synthesising enzymes was able to trigger branching in intact plants (Schluepmann *et al.*, 2003; Barbier *et al.*, 2015a). This suggests that HXK1 may be involved in promoting shoot branching when sugars are not readily available for buds due to the apical sugar sink strength, as it is the case in intact plants, while Tre6P is involved in triggering higher branching degree in response to increased sugar availability. However, it is possible that under our conditions the substrate of HXK1 (glucose) is limiting to increase HXK1 signalling activity compared with the wild-type, explaining why HXK1 over-expression could not increase shoot branching.

### HEXOKINASE1 interacts with cytokinin and strigolactone pathways to support shoot branching

Auxin and sugars act antagonistically to regulate bud outgrowth and fine-tune shoot architecture (Mason *et al.*, 2014; Dierck *et al.*, 2016; Schneider *et al.*, 2019; Barbier *et al.*, 2019b; Bertheloot *et al.*, 2020). The present study provides insight into the role of these signals in the regulation of apical dominance in arabidopsis. Our results demonstrate that the effect of auxin is largely dependent on sugar availability (Fig. 4-5). Indeed, in decapitated plants, we only observed a small effect of auxin on shoot branching, contrasting with a previous study in arabidopsis showing no effect of auxin on shoot branching in the wild-type Col-0 (Cline *et al.*, 2001). The inhibitory effect of auxin on shoot branching and bud outgrowth was stronger in defoliated plants and in the presence of low sucrose in the *in vitro* growth medium, respectively. This suggests that, similarly to pea and rose and chrysanthemum (Mason *et al.*, 2014; Dierck *et al.*, 2016; Bertheloot *et al.*, 2020), apical dominance in arabidopsis is maintained by an antagonistic action between auxin and sugars (Fig. 4-5).

The inhibitory effect of auxin on shoot branching was also dependent on the presence of HXK1 (Fig. 4-5). Additionally, the inhibitory effect of *hxk1* mutation on shoot branching was similar to the effect of defoliation or low sugar availability. This suggests that HXK1 is involved in sensing the leaf-derived sugar signal and in antagonizing the effect of auxin in decapitated plants.

Over-expressing HXK1 could not alleviate promote shoot branching in intact plants. Additionally, mannose, a sugar that triggers the HXK signalling pathway, could not alleviate the effect of auxin on bud outgrowth in rose, contrary to glucose (Supp. Fig. S7). This suggests that solely triggering the HXK signalling component is not enough to antagonize the inhibitory effect of auxin. However, in a similar manner to defoliation, HXK1 deficiency clearly led to increased sensitivity to auxin. This suggests that HXK1 may be involved in antagonizing the inhibitory effect of auxin when sugars are not readily available for buds.

The inhibitory effect of auxin on shoot branching is largely mediated by the antagonistic action of CK and SL (Rameau *et al.*, 2015; Walker & Bennett, 2018; Barbier *et al.*, 2019b). Sugars have been reported to promote CK synthesis and accumulation in rose stems and arabidopsis shoots (Barbier *et al.*, 2015b; Kiba *et al.*, 2019). Our results clearly demonstrate that *gin2* is CK-deficient and that the expression of CK-related genes, including the CK synthesis gene *IPT3*, are well correlated with *HXK1* expression under different sugar availability conditions. CK treatment partially restored the branching phenotype of *gin2*, further supporting the claim that the decreased branching phenotype of *gin2* is, at least partly, due to an impairment of the CK pathway. In arabidopsis, CK-deficient mutants have a decreased branching phenotype in intact plants, but not in response to decapitation, which increases sugar availability for buds (Müller *et al.*, 2015). We previously hypothesized that CK levels are important to promote bud outgrowth when sugars are not readily available for buds (Barbier *et al.*, 2019b). Accordingly, CK has been reported to promote nutrient sink strength (Roitsch & Ehneß, 2000; Werner *et al.*, 2008; Peleg *et al.*, 2011; Wang *et al.*, 2016), and it was recently reported that thus sugar-induced bud outgrowth in dark-grown potato sprouts was mediated, at least partly, by a CK-induced invertase activity (Salam *et al.*, 2020). In this view, we would expect that cytokinin synthesis mutants would indeed have a reduced response to decapitation which enhances sucrose availability. Our study shows a that the *gin2* mutant has a decreased branching phenotype mostly visible in intact plants. We suggest that the HXK1/CK pathway plays a role in maintaining shoot branching under nutrient restricting conditions such as is the case in buds of intact plants that are sugar-deprived due to the strong sink strength of the shoot apex (Mason *et al.*, 2014; Fichtner *et al.*, 2017).

The ability of cytokinins to promote branching in *gin2* also suggests a signaling role of HXK1, as the addition of cytokinins does not involve adding more carbon to the system. In the hypothesis mentioned above, the role of CK downstream of HXK1, would be to promote sugar sink strength within the axillary buds and to relocate carbon towards them. Similarly, this highly branched double mutant phenotype of *gin2* and SL mutants supports a signalling rather than catalytic role for HXK1.

High sugar availability has been reported to inhibit the effect of SL on bud outgrowth (Dierck *et al.*, 2016; Bertheloot *et al.*, 2020). We report here that the expression of *MAX2*, involved in SL-signalling, is negatively correlated with *HXK1* expression levels in response to sugar availability (Fig. 2 and Supp. Fig. S4). Additionally, inhibiting the SL pathway strongly induced shoot branching in the *gin2* background (Fig. 7), suggesting that a HXK1-dependent pathway may antagonize the perception of the SL pathway. Interestingly, glucose and SL have previously been found to interact during seedling development in arabidopsis (Li *et al.*, 2016), and sugar supply has been reported to alleviate the SL-induced leaf senescence in bamboo (Tian *et al.*, 2018). Altogether, these observations support the hypothesis that sugar and strigolactone pathways interact, and that these interactions may not be limited to the regulation of shoot branching (Fig. 8). Further studies are required to unravel the molecular mechanisms through which HXK1 interacts with these hormones to regulate shoot branching.

## CONCLUSIONS

The impact of sugar availability on shoot branching has been reported in different studies through manipulation of sugar supply or partitioning. This study highlights the involvement of a metabolic and signalling component of the sugar network in the control of shoot branching, namely HXK1. During this process, HXK1 senses the sugar status under restrictive conditions, allowing plants to fine-tune their architecture to adjust to their environment. Our results demonstrate that the HXK1-dependent pathway interacts with auxin, CK and SL to modulate shoot branching. This work should also prompt new studies to understand the role of Tre6P and other unexplored sugar signalling pathways involved in the control shoot branching.

## ACKNOWLEDGMENTS

We would like to thank Fengxi Han for her participation in the experiments performed in this study. We also would like to thank John Lunn, Jen Sheen, and Dani Eshel and Jessica Bertheloot for productive discussions, David Granot for plant material supply and Ottoline Leyser for helping with the genotyping of the *35S:HXK1-32* line. This research was funded by The University of Queensland and an Australian Research Council Discovery grant (DP150102086) and Laureate Fellowship (FL180100139).

## AUTHOR CONTRIBUTIONS

F. Barbier, C.A. Beveridge and S. Sakr and J. Le Gourrierec planned and designed the research.

F. Barbier, D. Cao, F. Fichtner, C. Weiste, M.D. Perez-Garcia and M. Caradeuc performed the experiments and analysed the data.

F. Barbier, D. Cao, F. Fichtner, C. Weiste, S. Sakr and C. Beveridge contributed to manuscript writing.

## SUPPORTING INFORMATION

**Supplemental Table 1.** Primers used in this study.

**Supplemental Table 2.** Statistical analysis of the Figure 6B.

**Supplemental Figure S1.** Pictures of arabidopsis buds and branching used for phenotypic analysis in this study.

**Supplemental Figure S2.** Impact of glucose analogues on bud elongation and *RMS4/MAX2* expression in pea buds.

**Supplemental Figure S3. A**. Impact of glucose analogues on the dry weight of single rose nodes.

**Supplemental Figure S4.** *HXK1* transcriptional response to stimuli controlling shoot branching.

**Supplemental Figure S5.** Length of the axillary rosette bud from Figure 4A divided by the rosette diameter

**Supplemental Figure S6.** Impact of night extension on early bud outgrowth of *max4-1*.

**Supplemental Figure S7.** Impact of glucose, mannose and auxin on bud elongation in rose.

## Notes

### Competing Interest Statement

The authors have declared no competing interest.

## REFERENCES

Barbier FF, Chabikwa TG, Ahsan MU, Cook SE, Powell R, Tanurdzic M, Beveridge CA. 2019a. A phenol/chloroform-free method to extract nucleic acids from recalcitrant, woody tropical species for gene expression and sequencing. Plant Methods 15: 62.

Barbier FF, Dun EA, Beveridge CA. 2017. Apical dominance. Current Biology 27: R864–R865.

Barbier FF, Dun EA, Kerr SC, Chabikwa TG, Beveridge CA. 2019b. An Update on the Signals Controlling Shoot Branching. Trends in Plant Science 24: 220–236.

Barbier FF, Lunn JE, Beveridge CA. 2015a. Ready, steady, go! A sugar hit starts the race to shoot branching. Current Opinion in Plant Biology 25: 39–45.

Barbier F, Péron T, Lecerf M, Perez-Garcia M-D, Barrière Q, Rolčík J, Boutet-Mercey S, Citerne S, Lemoine R, Porcheron B, et al. 2015b. Sucrose is an early modulator of the key hormonal mechanisms controlling bud outgrowth in *Rosa hybrida*. Journal of Experimental Botany 66: 2569–2582.

Bertheloot J, Barbier F, Boudon F, Perez-Garcia MD, Péron T, Citerne S, Dun E, Beveridge C, Godin C, Sakr S. 2020. Sugar availability suppresses the auxin-induced strigolactone pathway to promote bud outgrowth. New Phytologist 225: 866–879.

Brewer PB, Dun EA, Ferguson BJ, Rameau C, Beveridge CA. 2009. Strigolactone acts downstream of auxin to regulate bud outgrowth in pea and Arabidopsis. Plant Physiol. 150: 482–493.

Cao D, Lutz A, Hill CB, Callahan DL, Roessner U. 2016. A Quantitative Profiling Method of Phytohormones and Other Metabolites Applied to Barley Roots Subjected to Salinity Stress. Frontiers in Plant Science 7: 2070.

Chen J-G, Jones AM. 2004. AtRGS1 Function in Arabidopsis thaliana. Methods in Enzymology 389: 338–350.

Chevalier F, Nieminen K, Sánchez-Ferrero JC, Rodríguez ML, Chagoyen M, Hardtke CS, Cubas P. 2014. Strigolactone Promotes Degradation of DWARF14, an α/β Hydrolase Essential for Strigolactone Signaling in Arabidopsis. The Plant Cell 26: 1134–1150.

Cho Y-H, Yoo S-D, Sheen J. 2006. Regulatory functions of nuclear hexokinase1 complex in glucose signaling. Cell 127: 579–589.

Cline MG. 1996. Exogenous Auxin Effects on Lateral Bud Outgrowth in Decapitated Shoots. Annals of Botany 78: 255–266.

Cline MG, Chatfield SP, Leyser O. 2001. NAA Restores Apical Dominance in the axr3-1 Mutant of Arabidopsis thaliana. Annals of Botany 87: 61–65.

Coore HG, Randle PJ. 1964. Inhibition of glucose phosphorylation by mannoheptulose. Biochemical Journal 91: 56–59.

Cortès S, Gromova M, Evrard A, Roby C, Heyraud A, Rolin DB, Raymond P, Brouquisse RM. 2003. In Plants, 3-O-Methylglucose Is Phosphorylated by Hexokinase But Not Perceived as a Sugar. Plant Physiology 131: 824–837.

De la Fuente ASG, Villar-Palasí C, Asensio C. 1958. Substrate specificity and some other properties of baker’s yeast hexokinase. Biochimica et Biophysica Acta 30: 92–101.

Dierck R, Dhooghe E, Van Huylenbroeck J, De Riek J, De Keyser E, Van Der Straeten D. 2016. Response to strigolactone treatment in chrysanthemum axillary buds is influenced by auxin transport inhibition and sucrose availability. ACTA PHYSIOLOGIAE PLANTARUM.

Domagalska MA, Leyser O. 2011. Signal integration in the control of shoot branching. Nat. Rev. Mol. Cell Biol. 12: 211–221.

Evers JB. 2015. Sugar as a key component of the shoot branching regulation network. Plant, Cell & Environment 38: 1455–1456.

Evers JB, van der Krol AR, Vos J, Struik PC. 2011. Understanding shoot branching by modelling form and function. Trends in Plant Science 16: 464–467.

Feng J, Zhao S, Chen X, Wang W, Dong W, Chen J, Shen J-R, Liu L, Kuang T. 2015. Biochemical and structural study of Arabidopsis hexokinase 1. Acta Crystallographica. Section D, Biological Crystallography 71: 367–375.

Fichtner F, Barbier FF, Annunziata MG, Feil R, Olas JJ, Mueller-Roeber B, Stitt M, Beveridge CA, Lunn JE. In press. Regulation of shoot branching in Arabidopsis by trehalose 6-phosphate. New Phytologist DOI: 10.1111/nph.17006.

Fichtner F, Barbier FF, Feil R, Watanabe M, Annunziata MG, Chabikwa TG, Höfgen R, Stitt M, Beveridge CA, Lunn JE. 2017. Trehalose 6-phosphate is involved in triggering axillary bud outgrowth in garden pea (*Pisum sativum* L.). The Plant Journal: For Cell and Molecular Biology 92: 611–623.

Fichtner F, Lunn JE. 2021. The role of trehalose 6-phosphate (Tre6P) in plant metabolism and development. Annual Review of Plant Biology In press.

Fichtner F, Olas JJ, Feil R, Watanabe M, Krause U, Hoefgen R, Stitt M, Lunn JE. 2020. Functional Features of TREHALOSE-6-PHOSPHATE SYNTHASE1, an Essential Enzyme in Arabidopsis. The Plant Cell 32: 1949–1972.

Granot D, Kelly G, Stein O, David-Schwartz R. 2014. Substantial roles of hexokinase and fructokinase in the effects of sugars on plant physiology and development. Journal of Experimental Botany 65: 809–819.

Guo W, Chen L, Herrera-Estrella L, Cao D, Tran L-SP. 2020. Altering Plant Architecture to Improve Performance and Resistance. Trends in Plant Science 25: 1154–1170.

Hall SM, Hillman JR. 1975. Correlative inhibition of lateral bud growth in *Phaseolus vulgaris* L. timing of bud growth following decapitation. Planta 123: 137–143.

Hayward A, Stirnberg P, Beveridge C, Leyser O. 2009. Interactions between auxin and strigolactone in shoot branching control. Plant Physiol 151: 400–12.

Hei S, Liu Z, Huang A, She X. 2018. The regulator of G-protein signalling protein mediates D-glucose-induced stomatal closure via triggering hydrogen peroxide and nitric oxide production in Arabidopsis. Functional Plant Biology 45: 509.

Herold A, Lewis DH. 1977. Mannose and Green Plants: Occurrence, Physiology and Metabolism, and Use as a Tool to Study the Role of Orthophosphate. The New Phytologist 79: 1–40.

Hruz T, Laule O, Szabo G, Wessendorp F, Bleuler S, Oertle L, Widmayer P, Gruissem W, Zimmermann P. 2008. Genevestigator v3: a reference expression database for the meta-analysis of transcriptomes. Advances in Bioinformatics 2008: 420747.

Hu D-G, Sun C-H, Zhang Q-Y, An J-P, You C-X, Hao Y-J. 2016. Glucose Sensor MdHXK1 Phosphorylates and Stabilizes MdbHLH3 to Promote Anthocyanin Biosynthesis in Apple. PLoS genetics 12: e1006273.

Huang J-P, Tunc-Ozdemir M, Chang Y, Jones AM. 2015. Cooperative control between AtRGS1 and AtHXK1 in a WD40-repeat protein pathway in Arabidopsis thaliana. Frontiers in Plant Science 6: 851.

Huang L, Yu L-J, Zhang X, Fan B, Wang F-Z, Dai Y-S, Qi H, Zhou Y, Xie L-J, Xiao S. 2019. Autophagy regulates glucose-mediated root meristem activity by modulating ROS production in Arabidopsis. Autophagy 15: 407–422.

Ishikawa S, Maekawa M, Arite T, Onishi K, Takamure I, Kyozuka J. 2005. Suppression of Tiller Bud Activity in Tillering Dwarf Mutants of Rice. Plant and Cell Physiology 46: 79–86.

Kalliola M, Jakobson L, Davidsson P, Pennanen V, Waszczak C, Yarmolinsky D, Zamora O, Palva ET, Kariola T, Kollist H, et al. 2020. Differential role of MAX2 and strigolactones in pathogen, ozone, and stomatal responses. Plant Direct 4: e00206.

Karve R, Lauria M, Virnig A, Xia X, Rauh BL, Moore B d. 2010. Evolutionary Lineages and Functional Diversification of Plant Hexokinases. Molecular Plant 3: 334–346.

Karve A, Moore B d. 2009. Function of Arabidopsis hexokinase-like1 as a negative regulator of plant growth. Journal of Experimental Botany 60: 4137–4149.

Karve A, Rauh BL, Xia X, Kandasamy M, Meagher RB, Sheen J, Moore BD. 2008. Expression and evolutionary features of the hexokinase gene family in Arabidopsis. Planta 228: 411–425.

Kelly G, David-Schwartz R, Sade N, Moshelion M, Levi A, Alchanatis V, Granot D. 2012. The Pitfalls of Transgenic Selection and New Roles of AtHXK1: A High Level of AtHXK1 Expression Uncouples Hexokinase1-Dependent Sugar Signaling from Exogenous Sugar1[W]. Plant Physiology 159: 47–51.

Kiba T, Takebayashi Y, Kojima M, Sakakibara H. 2019. Sugar-induced de novo cytokinin biosynthesis contributes to Arabidopsis growth under elevated CO2. Scientific Reports 9: 1–15.

Kim Y-M, Heinzel N, Giese J-O, Koeber J, Melzer M, Rutten T, Wirén NV, Sonnewald U, Hajirezaei M-R. 2013. A dual role of tobacco hexokinase 1 in primary metabolism and sugar sensing. Plant, Cell & Environment 36: 1311–1327.

Lastdrager J, Hanson J, Smeekens S. 2014. Sugar signals and the control of plant growth and development. Journal of Experimental Botany 65: 799–807.

Leduc N, Roman H, Barbier F, Péron T, Huché-Thélier L, Lothier J, Demotes-Mainard S, Sakr S. 2014. Light Signaling in Bud Outgrowth and Branching in Plants. Plants 3: 223–250.

Lemoine R, La Camera S, Atanassova R, Dédaldéchamp F, Allario T, Pourtau N, Bonnemain J-L, Laloi M, Coutos-Thévenot P, Maurousset L, et al. 2013. Source-to-sink transport of sugar and regulation by environmental factors. Frontiers in Plant Science 4: 272.

Li Z, Ding Y, Xie L, Jian H, Gao Y, Yin J, Li J, Liu L. 2020. Regulation by sugar and hormone signaling of the growth of Brassica napus L. axillary buds at the transcriptome level. Plant Growth Regulation 90: 571–584.

Li GD, Pan LN, Jiang K, Takahashi I, Nakamura H, Xu YW, Asami T, Shen RF. 2016. Strigolactones are involved in sugar signaling to modulate early seedling development in *Arabidopsis*. Plant Biotechnology 33: 87–97.

Li L, Sheen J. 2016. Dynamic and diverse sugar signaling. Current Opinion in Plant Biology 33: 116–125.

Li Y, Xu S, Wang Z, He L, Xu K, Wang G. 2018. Glucose triggers stomatal closure mediated by basal signaling through HXK1 and PYR/RCAR receptors in Arabidopsis. Journal of Experimental Botany 69: 1471–1484.

Mason MG, Ross JJ, Babst BA, Wienclaw BN, Beveridge CA. 2014. Sugar demand, not auxin, is the initial regulator of apical dominance. Proceedings of the National Academy of Sciences 111: 6092–6097.

Moore B, Zhou L, Rolland F, Hall Q, Cheng W-H, Liu Y-X, Hwang I, Jones T, Sheen J. 2003. Role of the Arabidopsis glucose sensor HXK1 in nutrient, light, and hormonal signaling. Science (New York, N.Y.) 300: 332–336.

Moraes TA, Mengin V, Annunziata MG, Encke B, Krohn N, Höne M, Stitt M. 2019. Response of the Circadian Clock and Diel Starch Turnover to One Day of Low Light or Low CO2. Plant Physiology 179: 1457–1478.

Morris SE, Cox MCH, Ross JJ, Krisantini S, Beveridge CA. 2005. Auxin dynamics after decapitation are not correlated with the initial growth of axillary buds. Plant Physiol. 138: 1665–1672.

Müller D, Waldie T, Miyawaki K, To JP, Melnyk CW, Kieber JJ, Kakimoto T, Leyser O. 2015. Cytokinin is required for escape but not release from auxin mediated apical dominance. Plant J. 82: 874–86.

Nelson DC, Scaffidi A, Dun EA, Waters MT, Flematti GR, Dixon KW, Beveridge CA, Ghisalberti EL, Smith SM. 2011. F-box protein MAX2 has dual roles in karrikin and strigolactone signaling in Arabidopsis thaliana. Proceedings of the National Academy of Sciences of the United States of America 108: 8897–8902.

Otori K, Tamoi M, Tanabe N, Shigeoka S. 2017. Enhancements in sucrose biosynthesis capacity affect shoot branching in Arabidopsis. Bioscience, Biotechnology, and Biochemistry 81: 1470–1477.

Otori K, Tanabe N, Tamoi M, Shigeoka S. 2019. Sugar Transporter Protein 1 (STP1) contributes to regulation of the genes involved in shoot branching via carbon partitioning in Arabidopsis. Bioscience, Biotechnology, and Biochemistry 83: 472–481.

Patzke K, Prananingrum P, Klemens PAW, Trentmann O, Rodrigues CM, Keller I, Fernie AR, Geigenberger P, Bölter B, Lehmann M, et al. 2019. The Plastidic Sugar Transporter pSuT Influences Flowering and Affects Cold Responses. Plant Physiology 179: 569–587.

Pego JV, Weisbeek PJ, Smeekens SCM. 1999. Mannose Inhibits Arabidopsis Germination via a Hexokinase-Mediated Step. Plant Physiology 119: 1017–1024.

Peleg Z, Reguera M, Tumimbang E, Walia H, Blumwald E. 2011. Cytokinin-mediated source/sink modifications improve drought tolerance and increase grain yield in rice under water-stress. Plant Biotechnology Journal 9: 747–758.

Potter KC, Wang J, Schaller GE, Kieber JJ. 2018. Cytokinin modulates context-dependent chromatin accessibility through the type-B response regulators. Nature Plants 4: 1102.

Prasad TK, Li X, Abdel-Rahman AM, Hosokawa Z, Cloud NP, Lamotte CE, Cline MG. 1993. Does Auxin Play a Role in the Release of Apical Dominance by Shoot Inversion in *Ipomoea nil*? Annals of Botany 71: 223–229.

Rabot A, Henry C, Ben Baaziz K, Mortreau E, Azri W, Lothier J, Hamama L, Boummaza R, Leduc N, Pelleschi-Travier S, et al. 2012. Insight into the role of sugars in bud burst under light in the rose. Plant & Cell Physiology 53: 1068–1082.

Rameau C, Bertheloot J, Leduc N, Andrieu B, Foucher F, Sakr S. 2015. Multiple pathways regulate shoot branching. Frontiers in Plant Science 5: 741.

Rasmussen A, Mason MG, Cuyper CD, Brewer PB, Herold S, Agusti J, Geelen D, Greb T, Goormachtig S, Beeckman T, et al. 2012. Strigolactones Suppress Adventitious Rooting in Arabidopsis and Pea. Plant Physiology 158: 1976–1987.

Roitsch T, Ehneß R. 2000. Regulation of source/sink relations by cytokinins. Plant Growth Regulation 32: 359–367.

Rolland F, Baena-Gonzalez E, Sheen J. 2006. Sugar sensing and signaling in plants: conserved and novel mechanisms. Annual Review of Plant Biology 57: 675–709.

Sachs T, Thimann KV. 1964. Release of lateral buds from apical dominance. Nature 201: 939–940.

Sakr S, Wang M, Dédaldéchamp F, Perez-Garcia M-D, Ogé L, Hamama L, Atanassova R. 2018. The Sugar-Signaling Hub: Overview of Regulators and Interaction with the Hormonal and Metabolic Network. International Journal of Molecular Sciences 19: 2506.

Salam BB, Barbier F, Danieli R, Ziv C, Spíchal L, Teper-Bamnolker P, Jiang J, Ori N, Beveridge C, Eshel D. 2020. Sucrose promotes etiolated stem branching through activation of cytokinin accumulation followed by vacuolar invertase activity. bioRxiv: 2020.01.08.897009.

Salam BB, Malka SK, Zhu X, Gong H, Ziv C, Teper-Bamnolker P, Ori N, Jiang J, Eshel D. 2017. Etiolated Stem Branching Is a Result of Systemic Signaling Associated with Sucrose Level. Plant Physiology: pp.00995.2017.

Schluepmann H, Pellny T, van Dijken A, Smeekens S, Paul M. 2003. Trehalose 6-phosphate is indispensable for carbohydrate utilization and growth in *Arabidopsis thaliana*. Proceedings of the National Academy of Sciences of the United States of America 100: 6849–6854.

Schneider A, Godin C, Boudon F, Demotes-Mainard S, Sakr S, Bertheloot J. 2019. Light Regulation of Axillary Bud Outgrowth Along Plant Axes: An Overview of the Roles of Sugars and Hormones. Frontiers in Plant Science 10.

Schneider CA, Rasband WS, Eliceiri KW. 2012. NIH Image to ImageJ: 25 years of image analysis. Nature Methods 9: 671–675.

Smith AM, Stitt M. 2007. Coordination of carbon supply and plant growth. Plant, Cell & Environment 30: 1126–1149.

Soundappan I, Bennett T, Morffy N, Liang Y, Stanga JP, Abbas A, Leyser O, Nelson DC. 2015. SMAX1-LIKE/D53 Family Members Enable Distinct MAX2-Dependent Responses to Strigolactones and Karrikins in Arabidopsis. Plant Cell 27: 3143–59.

Stirnberg P, Furner IJ, Ottoline Leyser HM. 2007. MAX2 participates in an SCF complex which acts locally at the node to suppress shoot branching. The Plant Journal: For Cell and Molecular Biology 50: 80–94.

Tian M, Jiang K, Takahashi I, Li G. 2018. Strigolactone-induced senescence of a bamboo leaf in the dark is alleviated by exogenous sugar. Journal of Pesticide Science 43: 173–179.

Umehara M, Hanada A, Yoshida S, Akiyama K, Arite T, Takeda-Kamiya N, Magome H, Kamiya Y, Shirasu K, Yoneyama K, et al. 2008. Inhibition of shoot branching by new terpenoid plant hormones. Nature 455: 195–200.

Usadel B, Bläsing OE, Gibon Y, Retzlaff K, Höhne M, Günther M, Stitt M. 2008. Global Transcript Levels Respond to Small Changes of the Carbon Status during Progressive Exhaustion of Carbohydrates in Arabidopsis Rosettes. Plant Physiology 146: 1834–1861.

Van den Ende W. 2014. Sugars take a central position in plant growth, development and, stress responses. A focus on apical dominance. Frontiers in Plant Science 5.

Van Dingenen J, Vermeersch M, De Milde L, Hulsmans S, De Winne N, Van Leene J, Gonzalez N, Dhondt S, De Jaeger G, Rolland F, et al. 2019. The role of HEXOKINASE1 in Arabidopsis leaf growth. Plant Molecular Biology 99: 79–93.

Walker CH, Bennett T. 2018. Forbidden Fruit: Dominance Relationships and the Control of Shoot Architecture. Annual Plant Reviews 1: 1–38.

Wang L, Dong Q, Zhu Q, Tang N, Jia S, Xi C, Zhao H, Han S, Wang Y. 2017. Conformational Characteristics of Rice Hexokinase OsHXK7 as a Moonlighting Protein involved in Sugar Signalling and Metabolism. The Protein Journal 36: 249–256.

Wang W, Hao Q, Tian F, Li Q, Wang W. 2016. Cytokinin-Regulated Sucrose Metabolism in Stay-Green Wheat Phenotype. PLoS ONE 11: e0161351.

Wang B, Smith SM, Li J. 2018. Genetic Regulation of Shoot Architecture. Annual Review of Plant Biology 69: 437–468.

Waters MT, Gutjahr C, Bennett T, Nelson DC. 2017. Strigolactone Signaling and Evolution. Annual Review of Plant Biology 68: 291–322.

Werner T, Holst K, Pörs Y, Guivarc’h A, Mustroph A, Chriqui D, Grimm B, Schmülling T. 2008. Cytokinin deficiency causes distinct changes of sink and source parameters in tobacco shoots and roots. Journal of Experimental Botany 59: 2659–2672.

