## Supplemental information for "HEXOKINASE1 interferes with cytokinin synthesis and strigolactone perception during sugar-induced shoot branching"

**Supplemental Table 1.** Primers used in this study.

| Targeted gene | Purpose | Primer sequence |
|---|---|---|
| *HXK1* | *gin2-1* genotyping | Fwd: AGTTCAACAATCATTTGCAG<br>Rev: TGGCGCTCTTTGGGTAGGTT |
| *TUB3* | qRT-PCR | Fwd: TTCACAGCAAGCTTACGGAGGTCA<br>Rev: TGGTGGAGCCTTACAACGCTACTT |
| *ACT* | qRT-PCR | Fwd: AGTGGTCGTACAACCGGTATTGT<br>Rev1: GATGGCATGAGGAAGAGAGAAAC<br>Rev2: GAGGAAGAGCATACCCCTCGTA<br>Rev3: GAGGATAGCATGTGGAAGTGAGAA |
| *IPT1* | qRT-PCR | Fwd: CGGCGTCGTTTAAAGCAGTG<br>Rev: CCGCTTCACAATCTTTACGCTT |
| *IPT3* | qRT-PCR | Fwd: GGGTTCGTGTCTGAGAGAGTTGA<br>Rev: CGGAAATCCGATTGCTTTCTT |
| *IPT5* | qRT-PCR | Fwd: GTTGAACTTCCAAGGAAACATGG<br>Rev: CGGAAAACGAGTGGCTAGGT |
| *IPT7* | qRT-PCR | Fwd: ACCTAACGGCCACCCAGTATAG<br>Rev: CCCCGAGGAATGATTAACAAGT |
| *CKX1* | qRT-PCR | Fwd: ATCAGCGGTCAAGCATTCAAG<br>Rev: CGCTTCTCAGAACAGGTTACGAC |
| *CKX5* | qRT-PCR | Fwd: CGACGTTGTAACTGGGAAAGGA<br>Rev: AGCTGGTTCGAGAGAGATTCGTG |
| *CKX6* | qRT-PCR | Fwd: CCGACATGGACCACAGATCAG<br>Rev: CCGCGTTATGATGCCAAACT |
| *ARR5* | qRT-PCR | Fwd: TAAAACGCGCAAAGATCTGAGTTATCAG<br>Rev: GTCTCAAAAATTGTAAGCCGAAAGAATCAG |
| *ARR6* | qRT-PCR | Fwd: CCAAATCCTCCAACATTCGT<br>Rev: ACTCTGAGCAAACGCTCGAT |
| *ARR7* | qRT-PCR | Fwd: TGTTCTTGCCGTCGATGATA<br>Rev: TGGCATTGAGTAATCCGTCA |
| *ARR15* | qRT-PCR | Fwd: TTCCCGAGAGAAAACAATGG<br>Rev: CCCACTCTCAACAGTCGTCA |
| *D14* | qRT-PCR | Fwd: GTTGGTCACTCTGTTTCCGCTATG<br>Rev: GCGAAAATCCGATCAAGATAAGC |
| *MAX2* | qRT-PCR | Fwd: ATCAACGAGCTTCTCAGATCCCTA<br>Rev: GGAGGTAAGTCTTCAGTCCAGTGATAG |

| Tested group(s) | *p*-value |
|---|---|
| Genotype | 0.00086 |
| NAA | < 2.2e-16 |
| Sucrose | < 2.2e-16 |
| Sucrose:NAA | 2.773e-12 |
| NAA:Genotype | 0.076 |
| Sucrose:Genotype | 0.0099 |
| NAA:Sucrose:Genotype | 0.013 |

**Supplemental Table 2.** Statistical analysis of a two-way ANOVA performed on data presented in Figure 6.

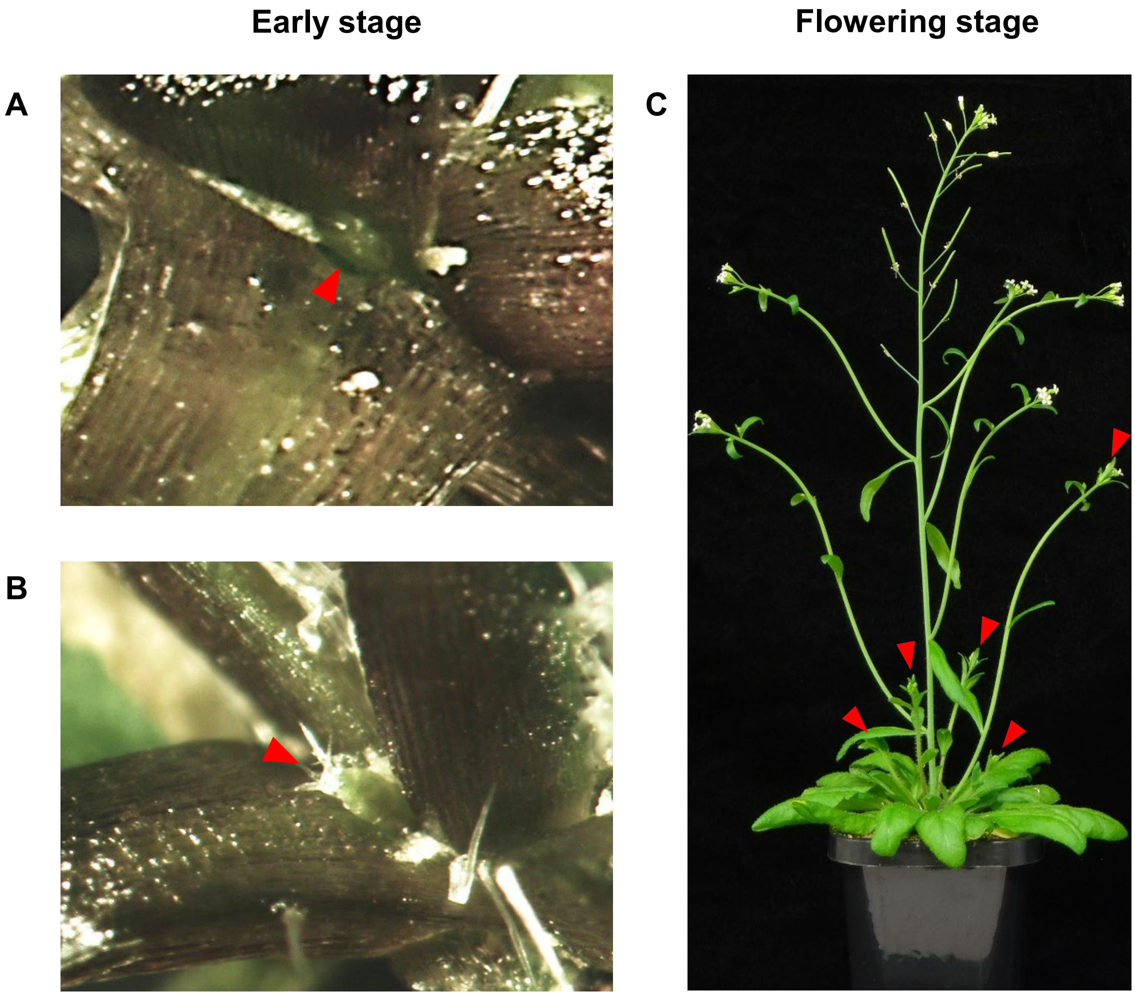


**Supplemental Figure S1. Pictures of arabidopsis buds and branching used for phenotypic analysis in this study.** Very young (A) and slightly older (B) axillary buds (red arrows) at the early stage (21 days after sowing). **C**. rosette branches (red arrows) at the flowering stage (38 days afters sowing)

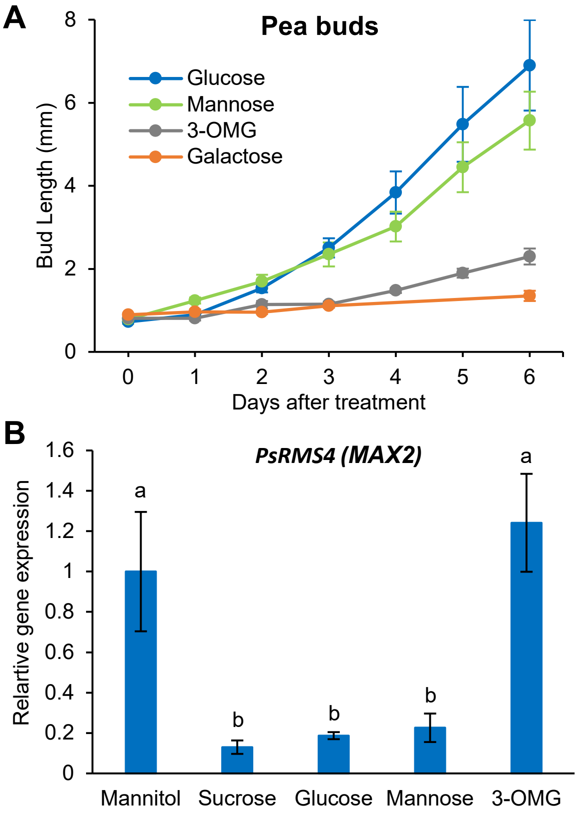


**Supplemental Figure S2. Impact of glucose analogues on bud elongation and *RMS4/MAX2* expression in pea buds. A**. Bud elongation in response to 30 mM glucose, mannose, 3 O-methylglucose (3-OMG) and galactose on single bud elongation *in vitro*. Data are mean ±se (n=8-10). **B**. *RMS4* expression pattern in pea buds fed with 30 mM mannitol, sucrose, glucose, mannose or 3-OMG. Buds were harvested from single node (node 5 and 6) excised from plants with five true leaves expanded and cultivated *in vitro*. Data are mean ±se (n=4 pools of 6 buds). Letters indicate significant difference between treatments (one-way ANOVA).

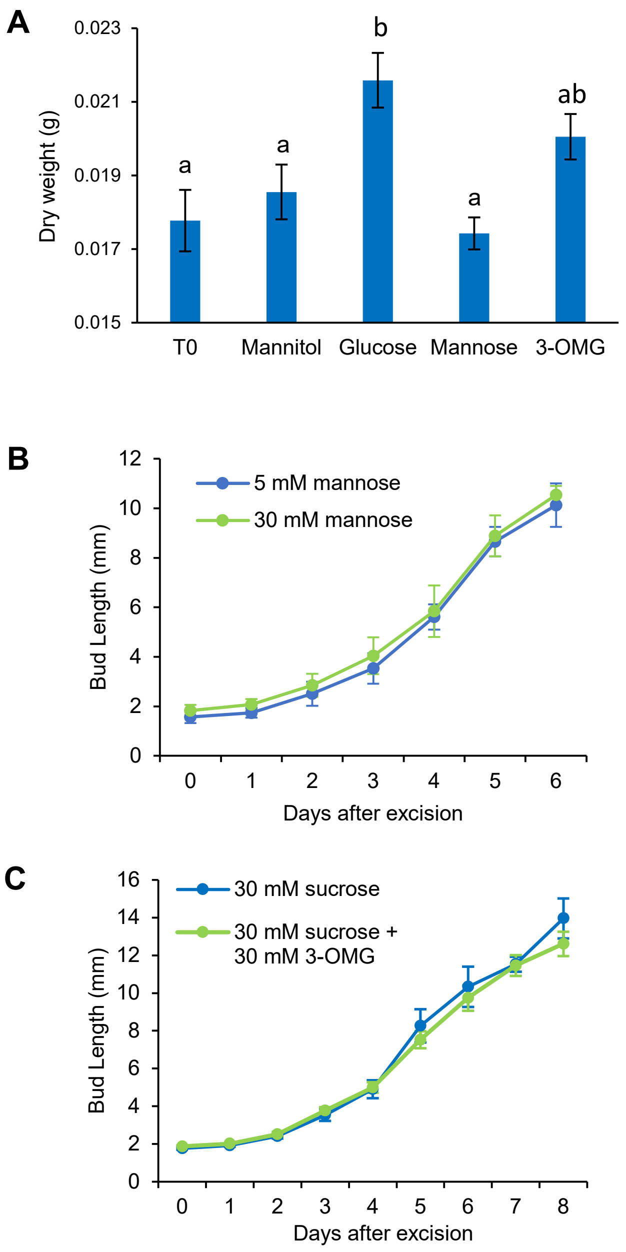


**Supplemental Figure S3. A**. Impact of 80 mM mannitol, glucose, mannose and 3-O-methylglucose (3-OMG) on the dry weight of single rose nodes (bud + stem) grown for three days**.** Data are mean ±se (n=10). Letters indicate significant difference between treatments (one-way ANOVA). **B**. Impact of 5 and 30 mM mannose on the elongation of rose buds grown *in vitro*. Data are mean ±se (n=10). **C**. Impact of 3-OMG on the sucrose induced bud outgrowth of single rose nodes. Data are mean ±se (n=10).

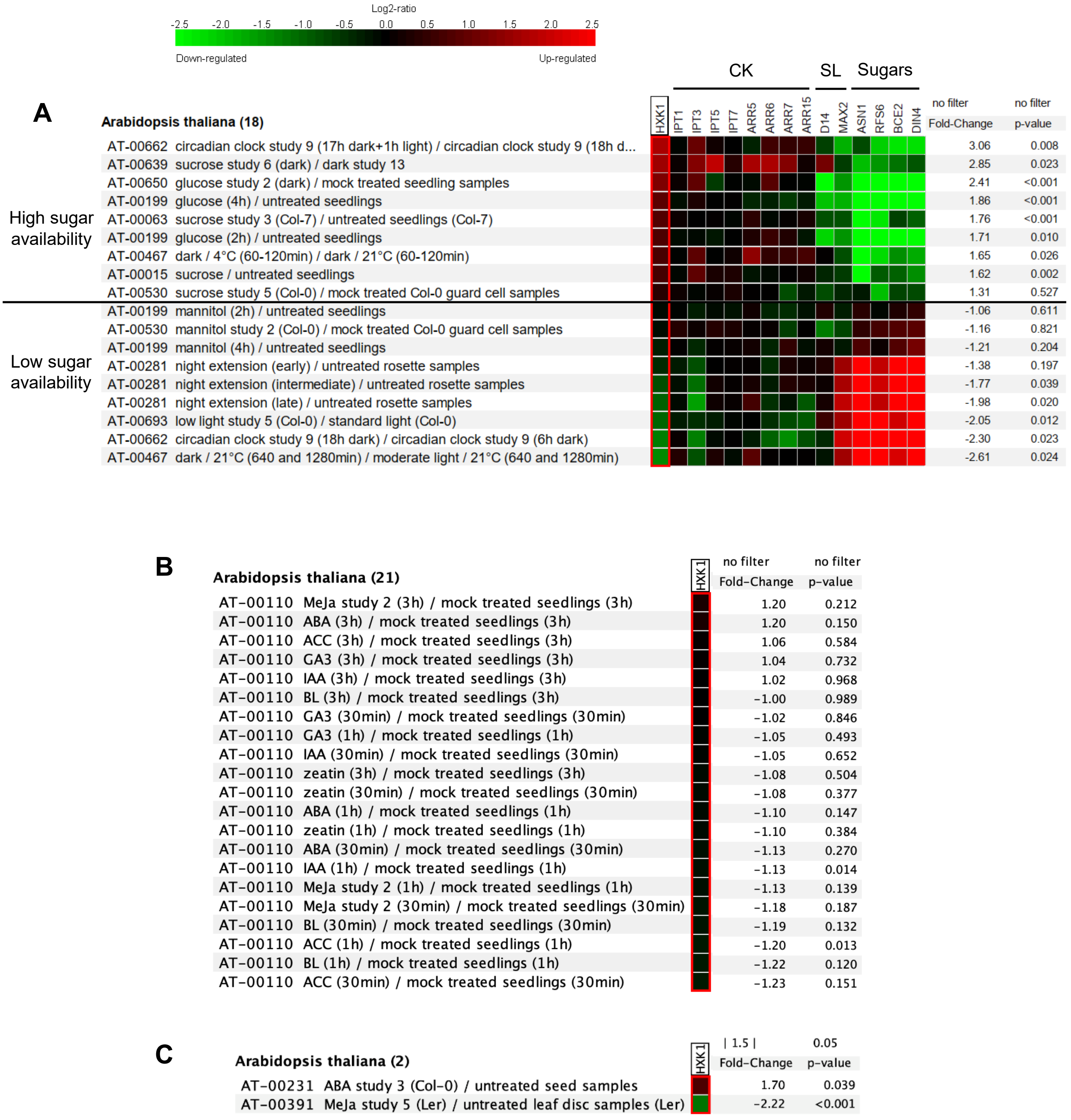


**Supplemental Figure S4. *HXK1* transcriptional response to stimuli controlling shoot branching. A**. Expression of *HXK1*, CK-related genes, SL-related genes and sugar markers (*ASN1* (*DIN6*, AT3G47340), *RS6* (*DIN10*, AT5G20250), *BCE2* (*DIN3*, AT3G06850) and *DIN4* (AT3G13450)) in different published transcriptomic analyses having in common the effect of to modulating sugar availability in plants as shown with the expression of the sugar marker genes. No filters have been set up. **B**. Expression of *AtHXK1* in arabidopsis seedlings treated for 30min, 1h and 3h with gibberellin (GA), jasmonic acid (MeJA0, ethylene precursor (ACC), cytokinin (BA), abscissic acid (ABA), auxin (IAA and zeatin) and brassinolid (BL). No filters have been set up. **C**. *HXK1* expression in the entire “hormones” dataset available in Genevestigator. The filters have been set up so that the displayed data have a *p*-value < 0.05 and a fold-change > 1.5.

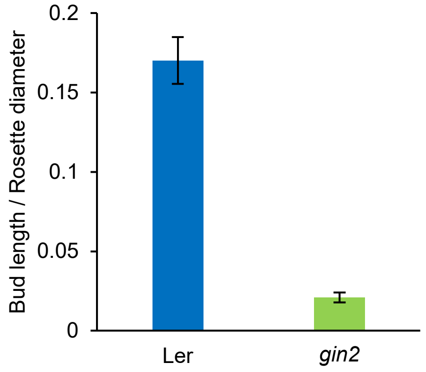


**Supplemental Figure S5.** Length of the axillary rosette bud (node 4) divided by the rosette diameter at an early stage in *gin2-1* and Landsberg *erecta* (Ler) 37 days after sowing. Data are mean ±se (n=15).

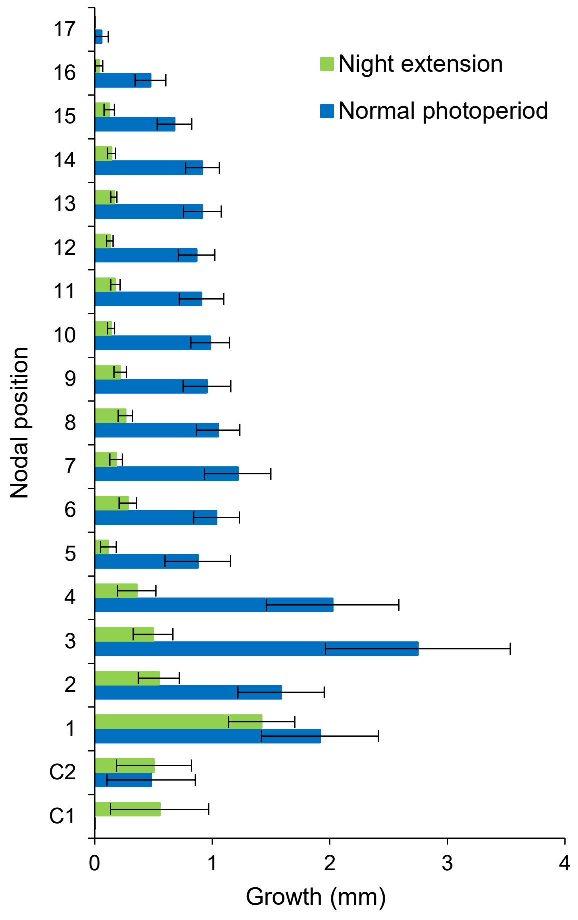


**Supplemental Figure S6. Impact of night extension on early bud outgrowth of *max4-1*.** Growth over 2 days of buds along the primary axis of 3 week old plants exposed to one night extension or to regular photoperiod (16h light, 8h night). Data are mean ±se (n=14-15).

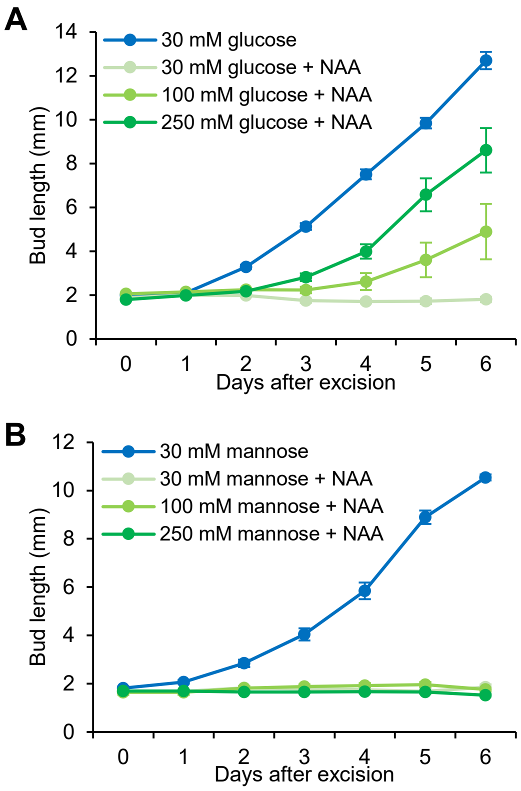


**Supplemental Figure S7. Impact of glucose, mannose and auxin (A,B) on bud elongation in rose.** Single nodes bearing a dormant bud were grown *in vitro* on media contaning 30, 100 and 250 mM glucose (**A**) or mannose (**B**) and either 0 or 2.5 µM NAA. Data are mean ±se (n=10).
